# Influenza A virus H1N1-derived circNP37 positively regulates viral replication by sponging host miR-361-5p

**DOI:** 10.1101/2023.09.04.556164

**Authors:** Chunyu Zhu, Jingyu Wang, Yalan Du, Chunli Li, Mengchan Hao, Yu Zhang, Xiaoqing Zhang, Yiwei Guan, Fangliang Zheng, Yuan Zhang, Jianjun Chen

## Abstract

RNA viruses, such as respiratory syncytial virus and SARS-CoV-2, can generate viral circular RNAs (circRNAs), which may play important roles during viral infection. However, whether influenza A viruses have this ability to generate viral circRNAs remains unknown. In this study, we discovered that the negative-strand RNA of the H1N1 nucleoprotein (NP) gene can generate a circRNA, designated circNP37. Furthermore, we demonstrated that circNP37 positively regulated viral replication by competitively sponging host miR-361-5p which inhibited polymerase basic protein 2 (PB2) expression. These results were confirmed using in vivo experiments. Compared with wild-type virus, infection with circNP37 knockout virus resulted in a reduced viral load in the lungs. This study demonstrates, for the first time, the existence and biological function of H1N1-derived circNP37. These findings help us better understand the mechanisms of influenza virus replication and pathogenicity.

**Highlights:**

- Negative-strand RNA of the H1N1 nucleoprotein gene can generate a circRNA
- CircNP37 plays important roles in viral replication and viral-host interactions
- CircNP37 positively regulates replication by competitively sponging host miR-361-5p

**eTOC Blurb:**

- In this study, we found that influenza A virus H1N1 infection can generate virus-derived circRNA, circNP37. We also demonstrated that circNP37 positively regulate viral PB2 gene expression and viral replication via sponge host miR-361-5p during viral infection.

## Introduction

Influenza A virus (IAV) is an enveloped, segmented, negative-stranded RNA virus.^1^ Historically, IAVs have been responsible for numerous large-scale epidemics.^2^ Because of antigenic drift and antigenic shift,^2^ which result in the constant generation of new viral strains, virus prevention and control are challenging. Therefore, IAV is a key concern for the protection of human public health.^3^

To better understand viral replication and identify new targets for treatment development, investigations are required at the viral protein and RNA levels.^4–6^ However, there are few studies on non-coding RNAs (ncRNAs) produced by IAV. ncRNAs operate as regulatory molecules at the RNA level, thus playing an important role in a wide range of biological processes.^7^ Using the classification of eukaryotic ncRNAs as a guide, ncRNAs can be further classified into housekeeping and regulatory ncRNAs. Housekeeping ncRNAs include ribosomal, transfer, and small nuclear RNAs. Regulatory ncRNAs include microRNAs (miRNAs), small RNAs (< 200 nt), long non-coding RNAs (> 200 nt), and circular RNAs (circRNAs).^8^

Several ncRNAs produced by IAV have been identified. For example, small viral RNAs (svRNAs)^9^ from H1N1 regulate the transition from transcription to replication by interacting with RNA dependent RNA polymerase (RdRP).^10^ Leader RNAs (leRNAs) are generated by H3N2 during infection.^11^ miR-HA-3p is produced by H5N1, which leads to increased virulence.^12^ These findings show that IAV-derived ncRNAs play a crucial role in viral replication and pathogenicity, which requires further investigation.

As a member of ncRNAs, circRNA is an RNA with a covalently closed circular structure^13^ that is abundant in archaea, prokaryotes, and eukaryotes.^14^ In addition to organisms with cellular structures, viruses can also produce circRNAs.^15^ These include human papillomavirus type 16,^16^ hepatitis B virus,^17^ Marek disease virus,^18^ Epstein-Barr virus (EBV),^19^ Kaposi’s sarcoma-associated herpesvirus,^20^ human cytomegalovirus,^21^ Merkel cell polyomavirus (MCV),^22^ and avian adenovirus type 4.^23^ Several studies have elucidated the biological functions and mechanisms of action of viral circRNAs. For example, the EBV-encoded circRNA LMP2A (ebv-circLMP2A) promotes cancer stem cells by binding to the miR-3908/ TRIM59/p53 axis. High expression of ebv-circLMP2A is significantly associated with metastasis and poor prognosis in patients with EBV-associated gastric cancer.^24^

Furthermore, MCV-derived circMCV-T regulates viral replication by interacting with MCV-derived miRNAs during viral infection.^25^

Currently, research on viral circRNAs is primarily focused on DNA viruses. Only a few RNA viruses have been reported to produce circRNAs, including respiratory syncytial virus,^26^ MERS-CoV,^27^ SARS-CoV-1,^27^ SARS-CoV-2,^27^ Bombyx mori cypovirus (BmCPV),^28^ murine hepatitis virus,^29^ and grass carp reovirus.^30^ Among the aforementioned RNA viruses that can produce circRNAs, only the function of the BmCPV circRNA has been revealed. It has been noted that BmCPV circRNA controls viral replication by influencing the ROS/NF-κB pathway through translational production of vSP27.^28^ However, whether viral circRNAs can be generated during influenza virus infection and the biological functions of circRNAs are unknown.

In this study, we found through in vitro and in vivo experiments that the IAV H1N1 (A/Puerto Rico/8/34, H1N1 [PR8]) generated a circRNA during infection, known as circNP37. Furthermore, we demonstrated that circNP37 competitively binds to the host miR-361-5p, which inhibits PB2 expression, thus positively regulating influenza virus PB2 expression and replication.

## Results

### Virus-derived circRNA can be generated during H1N1 infection

VirusCircBase is a database of circRNA molecules encoded by viruses and contains >46,000 circRNA predictions from 26 viral species.^31^ Six potential PR8-derived circRNAs with lengths greater than 200 nt predicted by VirusCircBase, IAV_circ_22, IAV_circ_30, IAV_circ_37, IAV_circ_38, IAV_circ_44, and IAV_circ_47 (Table S1), were chosen for further validation because of the high false-positive rates of circRNA predictions with less than 200 nt.^32^ Divergent primers were designed to verify these predicted circRNAs; however, only the primers designed for IAV_circ_37 amplified the back-splicing junction (BSJ) (Figures 1A, B, and S1). To determine the complete sequence of IAV_circ_37, two pairs of divergent primers were designed to amplify fragments with lengths close to the entire IAV_circ_37 sequence. PCR products with lengths of 313 and 367 bp were amplified (Figures 1A, C, and D), sequenced, and confirmed to be nucleoprotein (NP) negative-stranded RNAs with BSJ characteristics.

**Figure 1.**
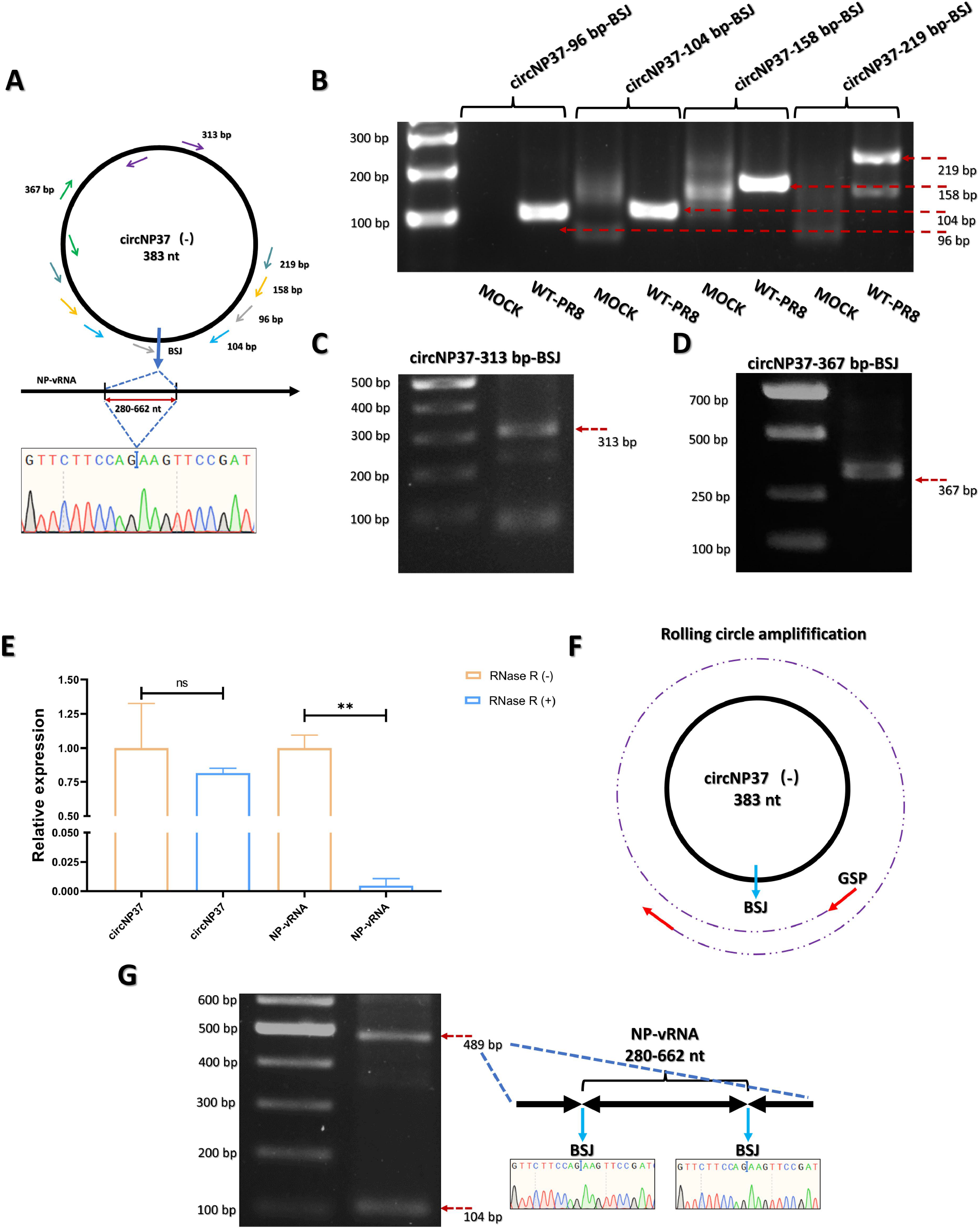
Virus-derived circRNA can be generated during H1N1 infection. (A) Schematic diagram of circNP37 divergent primers design. (B) RT-PCR results using several pairs of divergent primers validated by sequencing and confirmed as viral NP sequences with BSJ. (C and D) The 313 and 367 bp circNP37 sequences amplified using divergent primers after H1N1(PR8) infection. (E) RNA changes after RNase R digestion detected using real-time quantitative PCR. Statistical analysis performed using the unpaired Student’s *t* test. ns: not significant, \*\**p* < 0.01. (F and G) Validation of rolling circle amplification products. (F) Schematic diagram of circRNA rolling circle amplification. (G) A 489 bp fragment with rolling circle amplification characteristics discovered during RT-PCR using 104 bp BSJ amplification primers in numerous tests employing gene-specific primers (GSP) for reverse transcription.

The circular form of circRNAs make them resistant to digestion by nucleic acid exonucleases.^33^ RNase R digestion was performed to verify the IAV_circ_37. As shown in Figure 1E, the NP-vRNA sequence was significantly degraded after RNase R digestion, whereas the sequence of IAV_circ_37 did not change significantly.

Furthermore, due to their circular structure it is possible for circRNAs to produce a rolling circle amplification product when performing reverse transcription using a gene-specific primer (Figure 1F).

As shown in Figure 1G, primers that amplify a 104 bp fragment of IAV_circ_37 also amplify a 489 bp size product. Sequencing results showed that the 489 bp amplification product contained two BSJs and a complete IAV_circ_37 sequence between the two BSJs, indicating that the amplification product was the product of circNP37 rolling circle amplification.

These results indicated that the NP-negative-strand RNA of PR8 may produce circRNAs during viral infection. We designated it circNP37 according to the circRNA-naming principle proposed by Chen et al.^34^

### Knockout of circNP37 reduced viral transcription and replication

To elucidate the biological function of circNP37, a circNP37 knockout mutant virus was constructed. Since the circNP37 back-splice site (bss) flanks the conventional AG-GU base composition (Figure 2A),^35^ we hypothesized that the knockout of circNP37 without changing the NP protein sequence might be accomplished by circNP37 bss base mutation without affecting the amino acid coding of the viral native open reading frame (ORF). Thus, a pHW2000-NP plasmid with 5’ and 3’ bss mutations was constructed as shown in Figure 2A. After transfecting cells with wild-type pHW2000-NP, circNP37 was generated (Figure S2A). However, when cells were transfected with the 5’ + 3’ bss mutant pHW2000-NP, circNP37 could not be detected (Figure S2A).

**Figure 2.**
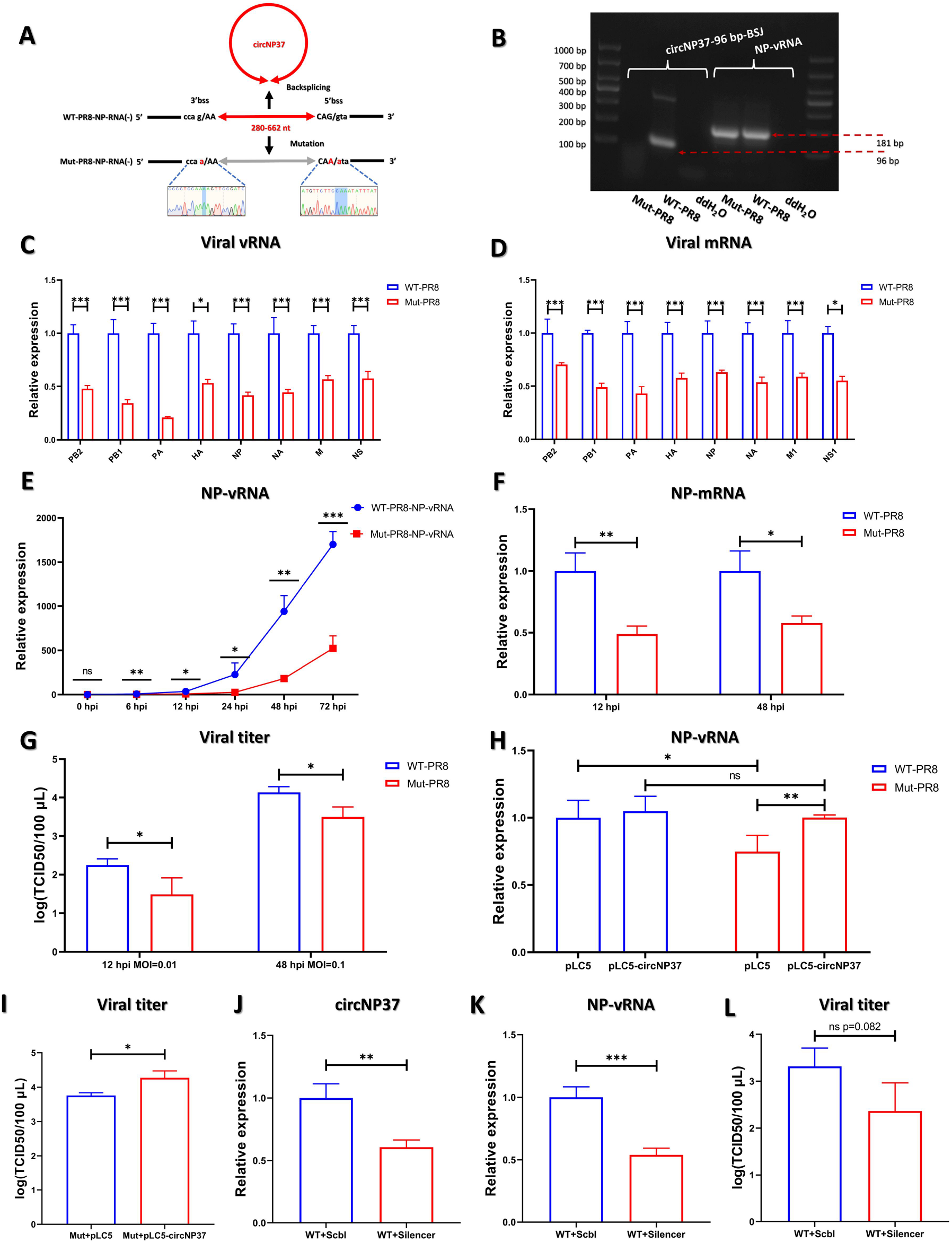
Knockout of viral circRNA reduces viral transcription and replication. (A) Schematic diagram of circNP37 back-splicing. In the WT-PR8-NP-RNA(-) diagram, red arrows with uppercase letters represent exons, lowercase letters with black lines represent introns, and the three linked bases represent a codon in the NP ORF. Red bases in Mut-PR8-NP-RNA(-) represent experimental changes in the bss. (B) Expression of viral NP-vRNA detected in both WT-PR8 and Mut-PR8 infected A549 cells, but circNP37 detected only after WT-PR8 infection. (C and D) Analysis of intracellular viral genome-wide replication and transcript levels following the infection of A549 cells with WT-PR8 or Mut-PR8 at 12 hpi (MOI = 0.01). (E) Differential alterations in the intracellular vRNA of the two viruses between WT-PR8 and Mut-PR8 infected A549 cells at MOI = 0.01 at various times. (F) Differential changes in intracellular viral mRNA levels at different time points in WT-PR8 and Mut-PR8 infected A549 cells at MOI = 0.01. (G) Differential variations in viral titers between WT-PR8 and Mut-PR8 infected A549 cells at MOI = 0.01 for 12 hpi or MOI = 0.1 for 48 hpi observed in cell supernatants. (H) Measurement of changes in intracellular vRNA in Mut-PR8 infected 293T cells overexpressing circNP37. (I) Measurement of changes in viral titers of Mut-PR8 infected 293T cells overexpressing circNP37. (J) Assessment of the knockdown effect of circNP37 by transfecting silencer targeting circNP37 into WT-PR8 infected 293T cells. (K-L) Measurement of variations in intracellular vRNA (K) and titer (L) of WT-PR8 infected 293T cells after circNP37 was specifically knocked down. Statistical analysis was performed using unpaired Student’s *t* test; ns: not significant, \**p* < 0.05, \*\**p* < 0.01, \*\*\**p* < 0.001.

We then used the 5’ + 3’ bss mutant pHW2000-NP to rescue cirNP37 knockout virus. We discovered that the bss mutation had no effect on virion assembly (Figures S2B and S2C) or viral infection (Figure 2B) but circNP37 could not be detected after viral infection (Figure 2B). We named the circNP37 knockout virus Mut-PR8 and the wild-type virus WT-PR8.

A549 cells were infected with WT-PR8 and Mut-PR8 to examine the effects of circNP37 on viral transcription and replication. At 12 hours post-infection (hpi), the total RNA of the infected cells was collected for quantitative analysis of the viral genome. Mut-PR8 had considerably lower levels of genome-wide transcription and replication than WT-PR8 (Figures 2C and 2D). To further study the effect of circNP37 on viral replication, A549 cells were infected with WT-PR8 or Mut-PR8 (MOI = 0.01), and tested for NP-vRNA expression at 0, 6, 12, 24, 48, and 72 hpi. As shown in Figure 2E, NP-vRNA levels in WT-PR8 infected cells were significantly higher than that in Mut-PR8 infected cells at all time points. Except for NP-vRNA, the transcription of NP-mRNA was also significantly decreased after Mut-PR8 infection compared to that after WT-PR8 infection at both 12 and 48 hpi (Figure 2F). Moreover, 12 and 48 h after viral infection (MOI = 0.01 and 0.1), the Mut-PR8 titer in the cell supernatant decreased compared to that of WT-PR8 (Figure 2G).

We restored circNP37 expression during Mut-PR8 infection using the circRNA-overexpression plasmid pLC5-circNP37. As shown in Figures 2H and 2I, when circNP37 was overexpressed during Mut-PR8 infection, NP-vRNA levels recovered to those of WT-PR8 infection, and viral titer was inc.

Next, we designed three different siRNA sequences and three different antisense oligonucleotide (ASO) sequences called silencer for circNP37 knockdown. 293T cells were infected with WT-PR8 8 h after transfection with a circNP37 silencer. At 36 hpi, circNP37 (Figure 2J), intracellular viral NP-vRNA levels (Figure 2K) and viral titer (Figure 2J) decreased. Combining these experimental findings, we found that circNP37 knockout/knockdown was detrimental to influenza virus replication, whereas circNP37 overexpression restored viral replication.

### CircNP37 is targeted and inhibited by miR-361-5p

The location of circNP37 in cells was identified to study its influence on viral replication. The proportion of circNP37 in the nucleus and cytoplasm of A549 cells was examined 24 h after WT-PR8 infection. The results showed that circNP37 was primarily localized in the cytoplasm (Figure 3A). Previous studies have shown that circRNAs can act as “miRNA sponges” in the cytoplasm^36^; therefore, we hypothesized that circNP37 may perform a biological function by sponging host miRNAs. Host miRNAs that could interact with circNP37 were predicted using miRanda (miRanda predictions were made using https://www.bioinformatics.com.cn, an online platform for data analysis and visualization) and miRDB (MicroRNA Target Prediction Database),^37^ using the human mature miRNA sequence information in miRbase^38^ as a reference. The intersection data of the two programs were then used to calculate the free energy of circNP37 binding to miRNA using RNAHybrid.^39^ Finally, miRmine^40^ was used to examine the candidate miRNAs for tissue and cell expression specificity, and three host miRNAs (miR-345-5p, miR-361-5p, and miR-769-3p) were screened that potentially interact with circNP37.

**Figure 3.**
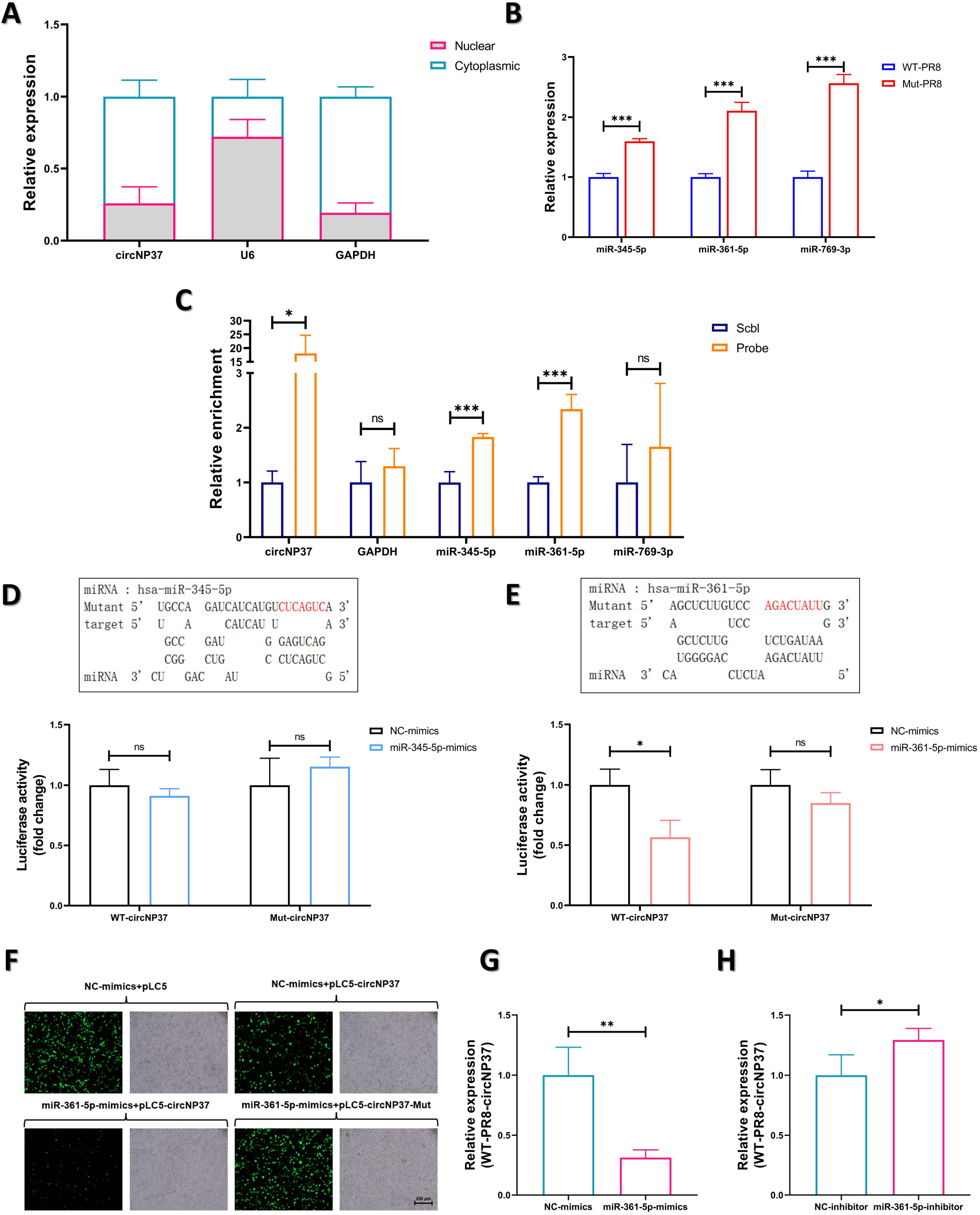
CircNP37 is targeted and inhibited by miR-361-5p. (A) Cellular location of circNP37 during viral infection after WT-PR8 infection of A549 cells determined by separating nuclear RNA from cytoplasmic RNA. (B) Determination of changes in expression levels of host miRNAs that might interact with circNP37 in A549 cells infected with WT-PR8 and Mut-PR8 at an MOI of 0.01 for 12 h. (C) CircNP37 and host miRNAs are significantly enriched in RAP experiments compared with scrambled probe (Scbl) pull-down experiments. (D and E) RNAHybrid prediction of circNP37 interaction with the miR-361-5p seed region, with the mutated seed region represented by red bases. A dual-luciferase experiment was used to detect the interaction between the miR-345-5p/miR-361-5p and circNP37 sequences. (F) Determination of the effect of simultaneous expression of miR-361-5p and pLC5-circNP37 on GFP expression in cells. Scale bar 250 μm. (G) Determination of the effect of miR-361-5p mimic overexpression on circNP37 expression in an infection model (WT-PR8-circNP37). (H) Determination of the effect of inhibition of miR-361-5p on circNP37 expression in infected cells (WT-PR8-circNP37). Error bars represent the mean ± SD. Statistical analysis: Unpaired Student’s *t* test; ns: not significant, \**p* < 0.05, \*\**p* < 0.01, \*\*\**p* < 0.001

A549 cells were infected with WT-PR8 and Mut-PR8 for 12 h, and differences in the expression of the three candidate miRNAs were detected. Compared to WT-PR8 infection, the expression of miR-345-5p, miR-361-5p, and miR-769-3p were higher in cells infected with Mut-PR8 (Figure 3B).

Biotin-labeled oligonucleotide probes with three different sequences targeting the circNP37 BSJ sequence were designed to confirm the interactions between circNP37 and these three host miRNAs. CircNP37 was overexpressed in 293T cells using pLC5-circNP37. After 48 h, RNA antisense purification (RAP) was performed using biotin-labeled probes to pull-down host miRNAs that could interact with circNP37. For miR-345-5p and miR-361-5p, the pull-down treatment group was significantly enriched, but not for miR-769-5p (Figure 3C), indicating that miR-345-5p and miR-361-5p may interact with circNP37.

The seed region for the interaction between circNP37 and miR-361-5p or miR-345-5p was predicted using RNAHybrid and miRanda. A dual-luciferase experiment was then performed to confirm this interaction. The full-length circNP37 sequence was cloned downstream of the firefly luciferase (Fluc) gene in pmiRGLO (Figure 3D). Fluc expression should decrease if miRNA targets the circNP37 sequence. As shown in Figure 3E, miR-361-5p considerably reduced Fluc fluorescence, whereas miR-345-5p did not (Figure 3D). Additionally, miR-361-5p was unable to effectively suppress Fluc expression when a mutation in the circNP37 seed region occurred (Figure 3E), confirming that miR-361-5p can interact with cirNP37.

Moreover, the circNP37 sequence was cloned upstream of the GFP gene in the circRNA expression vector (Figure S3A), and we anticipated that miR-361-5p might target the transcription product of the pLC5 expression vector through circNP37, which would then influence GFP expression. We co-transfected negative control (NC)-mimics/miR-361-5p-mimics and pLC5-circNP37/pLC5-circNP37-Mut (with a mutation in the interaction seed region) to further confirm the interaction between circNP37 and miR-361-5p. As shown in Figure 3F, cells transfected with NC-mimics+pLC5-circNP37 showed significantly lower GFP expression than those transfected with NC-mimics+pLC5, suggesting that the circNP37 sequence may be targeted by endogenous miR-361-5p in the cells. Further, the overexpression of miR-361-5p mimics further decreased the level of GFP expression. However, when the interaction seed region was mutated, the miR-361-5p mimics failed to decrease GFP expression, further confirming the interaction between miR-361-5p and circNP37 (Figure 3F).

Furthermore, when miR-361-5p was overexpressed or inhibited in 293T cells infected with WT-PR8, the expression level of circNP37 significantly decreased or increased, respectively (Figures 3G and 3H), providing evidence of the action of miR-361-5p on viral circRNA. Combined with the results of the previous experiment, it was shown that circNP37 could be targeted and inhibited by miR-361-5p.

### miR-361-5p targets PB2 and inhibits PB2 transcription and translation

CircNP37 and viral mRNA are primarily found in the cytoplasm during PR8 infection, and previous studies have shown that miR-361-5p functions in the cytoplasm.^41^ To investigate whether miR-361-5p plays a role in targeting viral mRNA, 8-gene mRNA of PR8 was expressed in 293T cells using the pCAGGS expression vector (Figures S3B and S3C). The cells were also transfected with biotin-labeled miR-361-5p and NC mimics (Figure S3D). As shown in Figure 4A, 48 h after transfection, miR-361-5p was found to significantly enrich the viral mRNA of the eight genes compared to the NC mimics by miRNA pull-down assay (PB2 and NA genes showed the most significant differences). This result indicated that miR-361-5p had the potential to interact with all eight PR8 genes and was most likely to interact with the PB2 and NA genes. As a component of the RdRp heterodimers, PB2 is critical for genome-wide transcription and replication.^35^ Since the presence of circNP37 previously influenced viral genome-wide transcription and replication (Figures 2A and 2B), we focused on the interactions among circNP37, miR-361-5p, and PB2.

**Figure 4.**
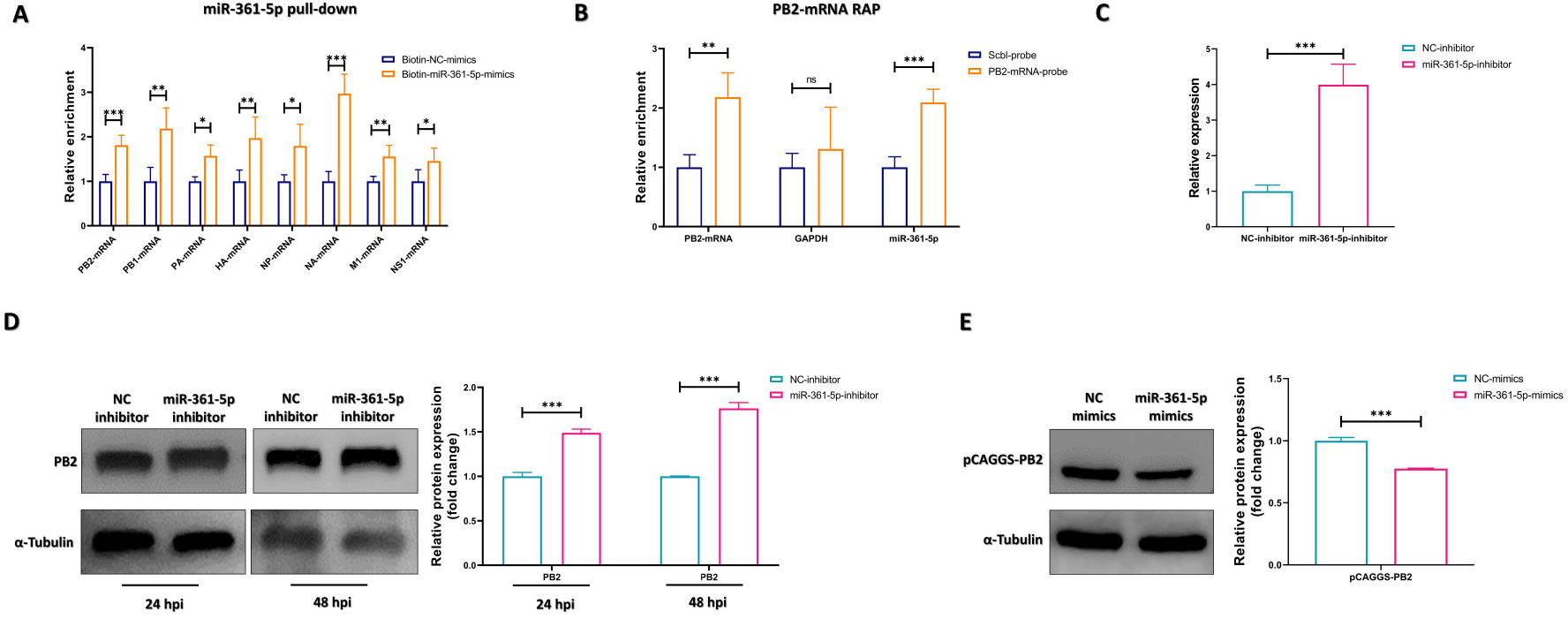
miR-361-5p targets PB2 and inhibits PB2 transcription and translation. (A) Pull-down examination of viral mRNA using biotin-labeled miR-361-5p demonstrating viral mRNA enrichment. (B) RAP experiments using biotin-labeled probes against PB2-mRNA with pull-down of PB2-mRNA to assess the enrichment of endogenous miR-361-5p in cells. (C) The effect on viral RNA level measured 36 hpi with Mut-PR8 (MOI = 0.01), with concomitant inhibition of miR-361-5p. (D) Determination of the effect of simultaneous inhibition of miR-361-5p during Mut-PR8 infection (MOI = 0.01) on the change in viral protein levels 36 hpi. (E) Determination of the effect on PB2 protein expression by overexpressing miR-361-5p while expressing PB2-mRNA in uninfected cells, and the effect on PB2 protein expression 24 hours post transfection. Protein expression was detected by grayscale analysis using blotting to determine the change in differential protein expression. Error bars represent the mean ± SD. Statistical analysis was performed using the unpaired Student’s *t* test, ns: not significant, \**p* < 0.05, \*\**p* < 0.01, \*\*\**p* < 0.001.

RAP experiments using probes targeting PB2-mRNA sequences to pull-down cellular endogenous miR-361-5p were carried out in PB2-mRNA overexpressed 293T cells. As shown in Figure 4B, miR-361-5p was significantly enriched by PB2-mRNA probes (Figure 4B). These results further confirmed the interaction between miR-361-5p and viral PB2-mRNA.

To study the influence of miR-361-5p on PB2 transcription and expression, we inhibited miR-361-5p during Mut-PR8 infection, which resulted in a significant increase in PB2 mRNA (Figure 4C) and protein expression (Figure 4D) levels compared to the NC inhibitor. To confirm the inhibitory effect of miR-361-5p on PB2 mRNA expression, we co-transfected miR-361-5p-mimics and PB2-mRNA in 293T cells. The results showed that miR-361-5p-mimics significantly reduced PB2 protein levels (Figure 4E), suggesting that miR-361-5p has the ability to target and inhibit PB2 expression.

### CircNP37 protects PB2 expression by sponging host miR-361-5p

We next studied the interactions among circNP37, miR-361-5p, and PB2-mRNA.

When cells were infected with Mut-PR8 lacking circNP37, overexpression of circNP37 by the pLC5-circNP37 plasmid resulted in a considerable increase in PB2 protein expression (Figure 5A). When cells were infected with WT-PR8 containing circNP37, overexpression of miR-361-5p considerably decreased PB2-mRNA (Figure 5B) and protein expression (Figure 5D) levels. However, when miR-361-5p was inhibited during WT-PR8 infection, the PB2-mRNA levels significantly increased (Figure 5C).

**Figure 5.**
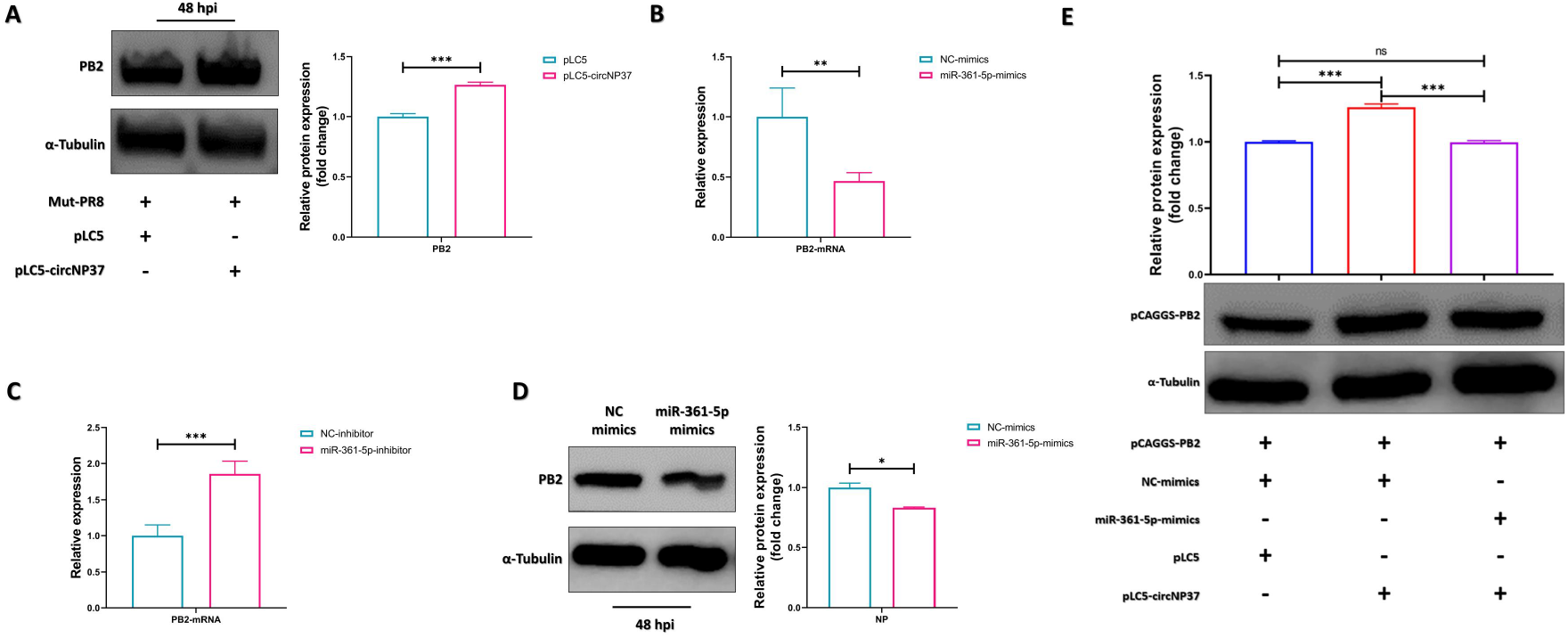
By sponging host miR-361-5p, circNP37 protects PB2 expression. (A) CircNP37 overexpression in Mut-PR8 infection promotes PB2 protein expression. (B) Transfection with miR-361-5p mimics has an inhibitory effect on PB2-mRNA expression in WT-PR8 cells. (C) Transfection of the miR-361-5p-inhibitor in WT-PR8 infection elevates PB2-mRNA expression. (D) Transfection of miR-361-5p-mimics significantly decreases PB2 protein expression in WT-PR8 infection. (E) The effect on PB2 protein expression was observed by controlling the expression of miR-361-5p and circNP37 under non-infected conditions. Grayscale analysis was used to determine protein expression from blots. Error bars represent the mean ± SD. Statistical analysis was performed using unpaired Student’s *t* test; ns: not significant, \**p* < 0.05, \*\**p* < 0.01, \*\*\**p* < 0.001.

To confirm the circNP37/miR-361-5p/PB2-mRNA interaction, 293T cells were transfected with pCAGGS-PB2, pLC5/pLC5-circNP37, or NC-mimics/miR-361-5p-mimics. As shown in Figure 5E, circNP37 alone resulted in significantly higher levels of PB2 protein expression (lane 2); however, when miR-361-5p was overexpressed along with circNP37, PB2 protein levels decreased (lane 3).

Taken together, we discovered that circNP37 reversed the negative effects of miR-361-5p on PB2 protein production in both infected and non-infected settings. It is possible that miR-361-5p inhibits PB2 mRNA by binding to it; however, circNP37 competitively binds miR-361-5p in this process, thus decreasing its inhibitory effect on PB2 mRNA.

### In vivo studies showed that circNP37 knockout inhibits viral replication

In vivo experiments were performed to confirm the effect of circNP37 on viral replication. Female BALB/c mice aged 5–7 weeks were intranasally infected with PBS, WT-PR8, or Mut-PR8. As shown in Figure 6A, circNP37 was only detected in the lungs of WT-PR8 infected mice. However, both WT-PR8 and Mut-PR8 were able to infect the mice in this study (Figure 6A). At 3 dpi, both groups displayed telltale symptoms of influenza infection, such as hunched posture, lethargy, labored breathing, and ruffled fur. Although variations in body weight (Figure 6B) and lung index (Figure 6D) between the two groups were not significant, WT-PR8-infected mice reached the endpoint (all deaths) faster than Mut-PR8-infected mice (Figure 6C).

**Figure 6.**
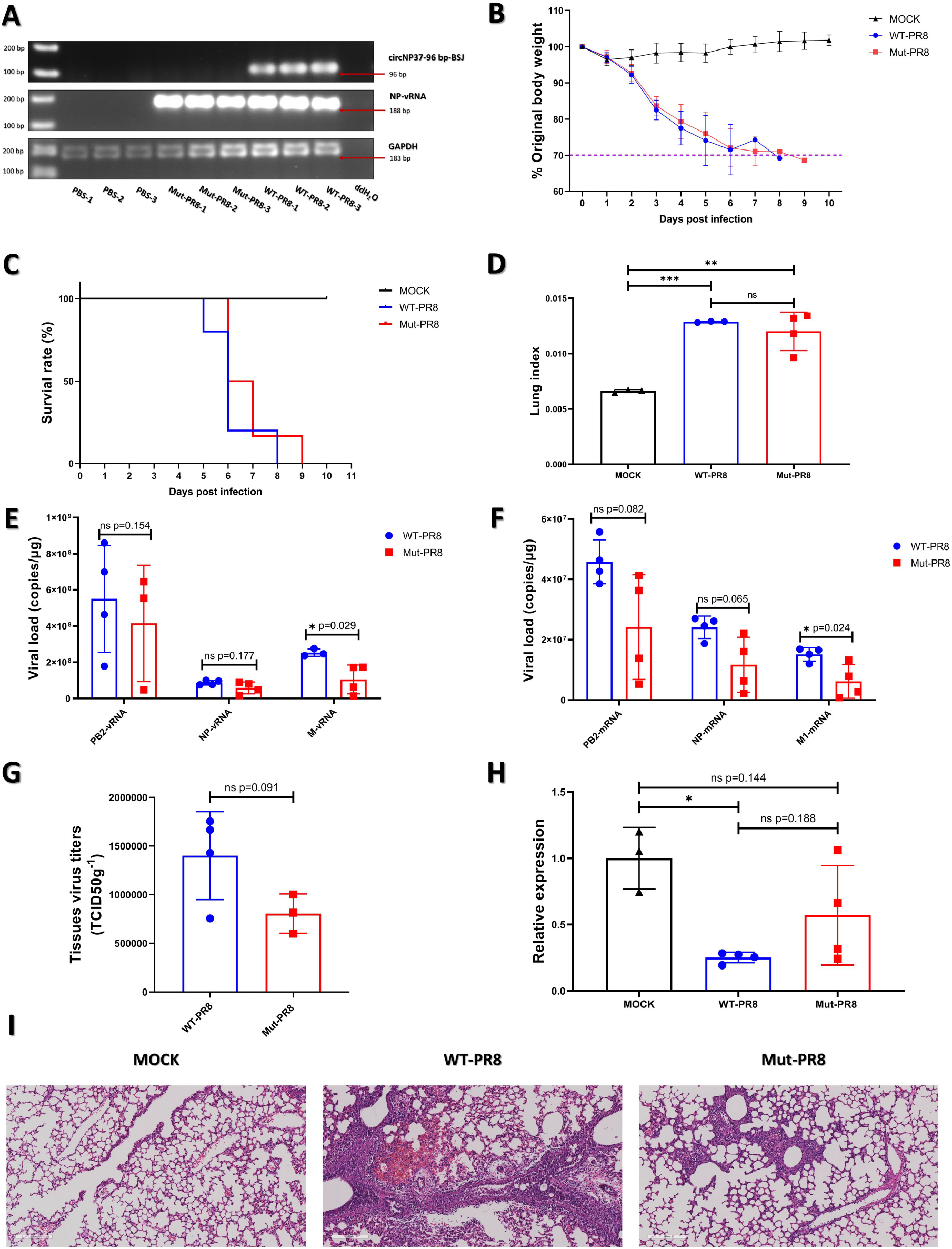
Knockout of circNP37 in in vivo experiments is detrimental to virus replication. (A) Lung circNP37 expression assay in WT-PR8 and Mut-PR8 infected mice. (B) Body weight changes in mice after WT-PR8 or Mut-PR8 infections (WT-PR8, n = 10; Mut-PR8, n = 11). (C) Differences in mouse survival following infection with WT-PR8 or Mut-PR8 (WT-PR8, n = 10, Mut-PR8: n = 11). (D) Differences in mouse lung indexes after infection with WT-PR8 or Mut-PR8 (n = 4); error bars indicate the mean ± SD. Survival analysis was performed using the log-rank (Mantel–Cox) test; the difference between the two treatment groups was compared in parallel using unpaired Student’s *t* test. (E) Lung PB2, NP, and M vRNA copy number assays after WT-PR8 or Mut-PR8 infection (n = 4). (F) PB2, NP, and M1 mRNA copy number assays after WT-PR8 or Mut-PR8 infection (n = 4). (G) Detection of viral titers in the lungs after WT-PR8 or Mut-PR8 infection (n = 4). (H) Detection of mmu-miR-361-5p in the lungs following infection with WT-PR8 compared with Mut-PR8 (n = 4). Error bars represent the mean ± SD. Statistical analysis was performed using an unpaired Student’s *t* test; ns: not significant, \**p* < 0.05, \*\**p* < 0.01, \*\*\**p* < 0.001. (I) Congestion, hemorrhage, serous effusion, lymphocytic infiltration, bronchodilatation, and alveolar rupture of the lungs (200 μM) of MOCK (PBS-treated mice), WT-PR8-infected, and Mut-PR8-infected mice.

Next, the viral loads in the lungs of WT-PR8 and Mut-PR8 infected mice were detected. Compared to WT-PR8 infected mice, PB2, NP, and M vRNA levels in the lungs of Mut-PR8 infected mice showed a decrease (Figure 6E). Moreover, the PB2, NP, and M1 mRNA levels in the lungs of Mut-PR8 infected mice were lower than that of WT-PR8 infected mice, with M1 mRNA showing a significant decrease (*p* = 0.024) and PB2 mRNA and NP mRNA exhibiting almost significant decreases (*p* = 0.082 and *p* = 0.065, respectively) (Figure 6F). In addition, the viral titer in the lungs of mice after infection with Mut-PR8 was lower than in the WT-PR8 infected group, nearing a significant level (*p* = 0.091) (Figure 6G).

miR-361-5p is sequence conserved across multiple species.^38^ We simultaneously found that the miR-361-5p level was lower in the lungs of WT-PR8 infected mice than in the PBS-treated group, and there was a trend toward an increase in miR-361-5p in the lungs of Mut-PR8 infected mice when compared to the WT-PR8 infected group; however, this difference was not statistically significant (*p* = 0.188) (Figure 6H). Taken together, these findings indicated that circNP37 knockout results in lower viral replication levels in vivo. CircNP37 knockout had the same effect on host miR-361-5p expression at the cellular level (Figure 3B).

Pathological sections were prepared from the left lungs of the mice at 3 dpi. As shown in Figure 6I, mice infected with WT-PR8 experienced more severe pathological alterations than mice infected with Mut-PR8, including congestion, hemorrhage, serous effusion, lymphocytic infiltration, bronchodilation, and alveolar rupture. These studies showed that in vivo experiments using circNP37 knockout mice somewhat reduced lung damage.

## Discussion

In this study, we demonstrate for the first time that IAV circRNAs are present during viral infection. These results showed that circNP37 positively regulates PR8 replication by competitively sponging host miR-361-5p to protect the PB2-mRNA during viral infection.

As previous studies have reported, viral circRNAs can regulate their own replication at the RNA level in various ways, including by interacting with host factors (e.g., EBV-derived circBART2.2^42^ and circLMP2A^24^) and becoming a part of viral-host interactions at the RNA level (e.g., MCV-derived circMCV-T^25^ and circALTOs^22^). For example, a previous report showed that the expression of SARS-CoV-2 derived circ_3205 was positively correlated with the expression of spike mRNA and the viral load.^43^ It has also been suggested that circ_3205 controls the expression of the host genes KCNMB4 and PRKCE by sponging hsa-miR-298.^43^ All these researches and our study demonstrated that virus-derived circRNA could serve as a “tool” for virus-host interaction.

To date, the exact mechanism by which RNA viral circRNAs are generated is unknown. This study suggests that the production of circNP37 might not be dependent on the process of viral infection because cells transfected with the pHW2000-NP plasmid could also generate circNP37 (Figure S2, A). According to Starke et al., conventional splicing elements are required to produce exonic circRNA in eukaryotes.^44^ Our research also confirmed that the AG-GU splice site is necessary for circNP37 production. As IAV undergoes transcriptional replication in the nucleus, NP-negative-strand RNA may interact with the intranuclear spliceosome to produce circRNAs during viral infection. However, respiratory syncytial virus, MERS-CoV, SARS-CoV-1, SARS-CoV-2, BmCPV, murine hepatitis virus, and grass carp reovirus are localized in the cytoplasm for transcriptional replication.^45^ How these viruses come into contact with the spliceosome in the nucleus during the infection process and whether they employ self-expressed proteins for circRNA generation requires further investigation.

Currently, RNA viral circRNA research is still at an early stage, and further work is needed to explain the mechanism of viral circRNA generation. For example, whether there is a specific class of splice sites that promote splicing, and the presence of other types of regulatory elements (such as flanking reverse complementary sequences and splicing enhancers) in RNA virus circRNA production remains to be determined. Investigating the mechanisms of viral circRNA generation can help expand our understanding of the viral life cycle and explore its applications. For example, investigation of viral circRNA generation has enabled the construction of engineered circRNA expression vectors that require the class I catalytic intron of the Td gene of the T4 phage that circulates circRNAs^46^ and the ribosomal entry site carried by the encephalomyocarditis virus as important elements.^46^ This enables the production of circRNA vaccines. Research has shown that in contrast to the mRNA vaccine, the circRNA vaccine of SARS-CoV-2 yields higher and longer-lasting antigen production and offers excellent protection^47^ owing to the stability conveyed by its circular structure.^47^ These studies imply that studies on the RNA virus circRNA generation mechanism may be useful not only theoretically, but also practically.^47^

Eukaryotic circRNAs play a fine-tuning role in a range of biological models.^48,49^ In this study, we discovered that the effect of circNP37 on viral replication can likewise be a method of fine-tuning. At the cellular level, circNP37 resulted in a 2–4-fold difference in viral RNA, protein, and titer levels when compared to the control treatment. The results of the in vivo experiments were consistent with those of the cell experiments. Furthermore, the lack of circNP37 resulted in considerably lower lung inflammation in mice. However, the presence or absence of circNP37 had no influence on body weight change, survival curve, or lung index in mice. These results indicate that circNP37 plays a “fine-tuning” role at the RNA level in virus infection and has some effects on viral load and titer. However, this is not crucial for viral pathogenicity in mice.

We also attempted to determine whether the circNP37/miR-361-5p interaction plays any other role during viral infection. The biological function of miR-361-5p during IAV infection has not been studied; however, it has been demonstrated that hsa-miR-361-5p is considerably downregulated during infection with influenza A viruses (H1N1 and H7N7).^50^ The biological function of hsa-miR-361-5p has also been investigated in cancer research.^51,52^ A previous report showed that TNF receptor-associated factor 3 (TRAF3) was targeted and inhibited by miR-361-5p.^53^ Since TRAF3 plays an important role in the NF-κB pathway,^54^ we also investigated its differential transcriptional changes in WT-PR8 and Mut-PR8 infections. TRAF3 transcriptional expression was significantly decreased in Mut-PR8-infected cells (data not shown). However, it remains unclear whether circNP37 causes significant variation in NF-κB pathway activation during viral infection, which requires further investigation.

In summary, this study demonstrated the existence and biological function of the H1N1-derived circRNA, circNP37, through in vitro and in vivo experiments. The working mechanism was also investigated. The results of this study provide evidence that this viral circRNA of H1N1 plays important roles in viral replication and viral-host interactions.

## Supporting information

Supplemental Iformation

## Acknowledgments

This work was supported by the National Natural Science Foundation of China (31970174, 32270180, 31600128) and the program of Liaoning Provincial Department of Education (LJC201910, LJKFZ20220179). We would like to thank Juan Min at the Centre for Instrumental Analysis and Metrology, Wuhan Institute of Virology Chinese Academy of Sciences, for providing technical assistance. We are grateful to He Zhao from the Center for Animal Experiment, Wuhan Institute of Virology Chinese Academy of Sciences, for help in analysis of pathological sections. Special thanks to Hanzhong Wang of Wuhan Institute of Virology, Chinese Academy of Sciences, for his contribution in this project.

## Author contributions

Chunyu Zhu, Jianjun Chen, Yuan Zhang, Fangliang Zheng and Jingyu Wang conceived and designed the projects. Jingyu Wang and Chunyu Zhu performed most of the experiments. Mengchan Hao and Yiwei Guan contributed to in vivo experiments. Yalan Du and Chunli Li contributed to RT-qPCR experiments. Xiaoqing Zhang and Yu Zhang contributed to virus titer determination. Jianjun Chen, Yuan Zhang, Fangliang Zheng and Jingyu Wang analyzed the data. Jianjun Chen, Yuan Zhang, Fangliang Zheng, Chunyu Zhu and Jingyu Wang wrote and edited the manuscript. All authors read and approved the final manuscript.

