## Supplemental Iformation for "Influenza A virus H1N1-derived circNP37 positively regulates viral replication by sponging host miR-361-5p"

**Supplemental information titles and legends**

**Table S1**

| | circRNA name | circRNA_start | circRNA_end | strand | gene | Junction read counts | circRNA_length |
|---|---|---|---|---|---|---|---|
| 1 | IAV_circ_22 | 325 | 1361 | + | HA | 16 | 1037 |
| 2 | IAV_circ_30 | 368 | 1411 | - | NA | 16 | 1044 |
| **3** | **IAV_circ_37** | **280** | **662** | **-** | **NP** | **33** | **383** |
| 4 | IAV_circ_38 | 307 | 1393 | + | NP | 65 | 1087 |
| 5 | IAV_circ_44 | 159 | 2148 | - | PB1 | 41 | 1990 |
| 6 | IAV_circ_47 | 1560 | 2028 | + | PA | 10 | 469 |

Table S1: Predictive information from VirusCircBase for viral circRNAs that may be generated in influenza A virus infections.

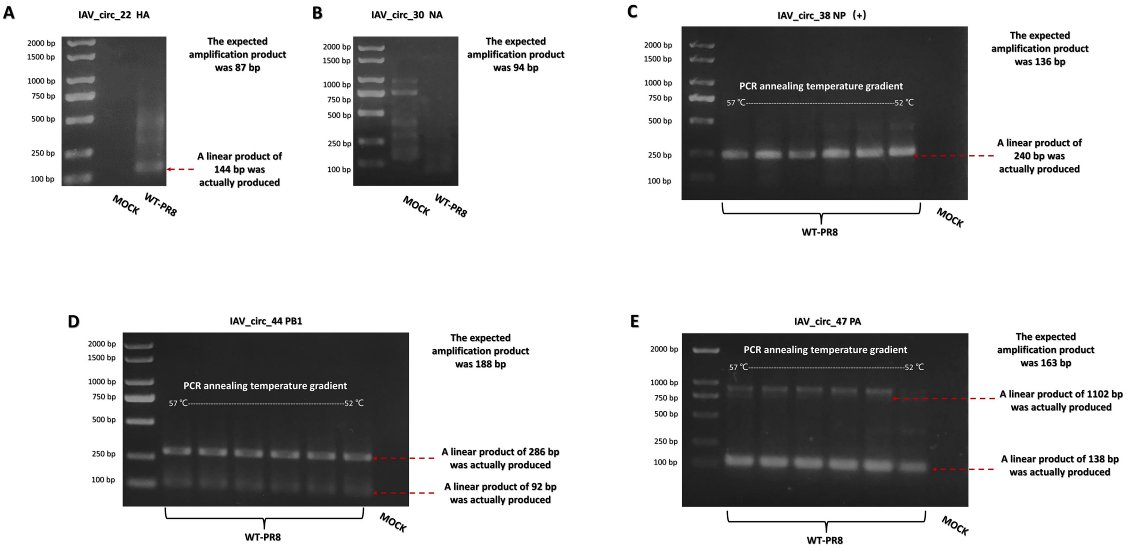


Figure S1. Predicted results from HA (A), NP (+) (B), NA (C), PB1 (D), and PA failed to validate the presence of viral circRNA

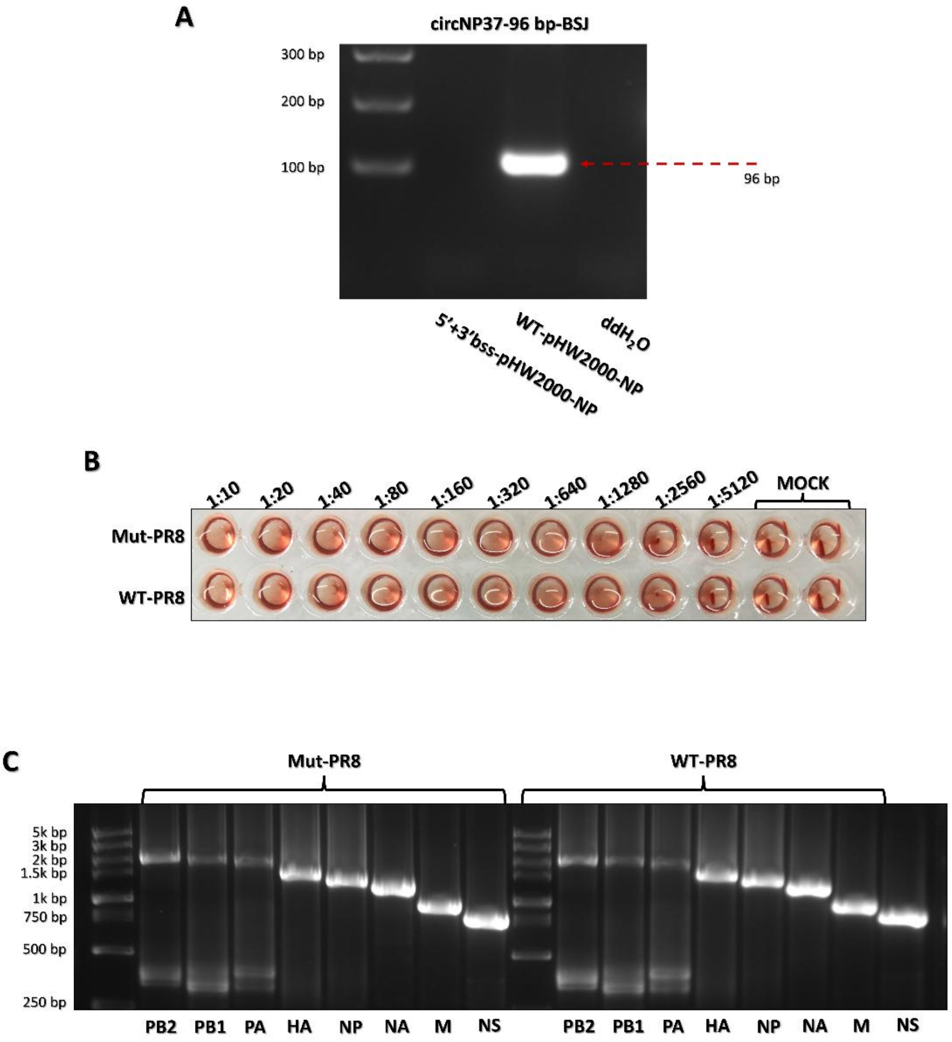


Figure S2. Rescue of circNP37 knockout PR8

A: CircNP37 was detectable when cells were transfected with WT-pHW2000-NP; however, after cells were transfected with pHW2000-NP with the 5'+3' bss mutation, circNP37 expression could not be found.

B: Rescue virus hemagglutination titer assay, 1:1280.

C: Genomic RNA RT-PCR detection of rescue viruses.

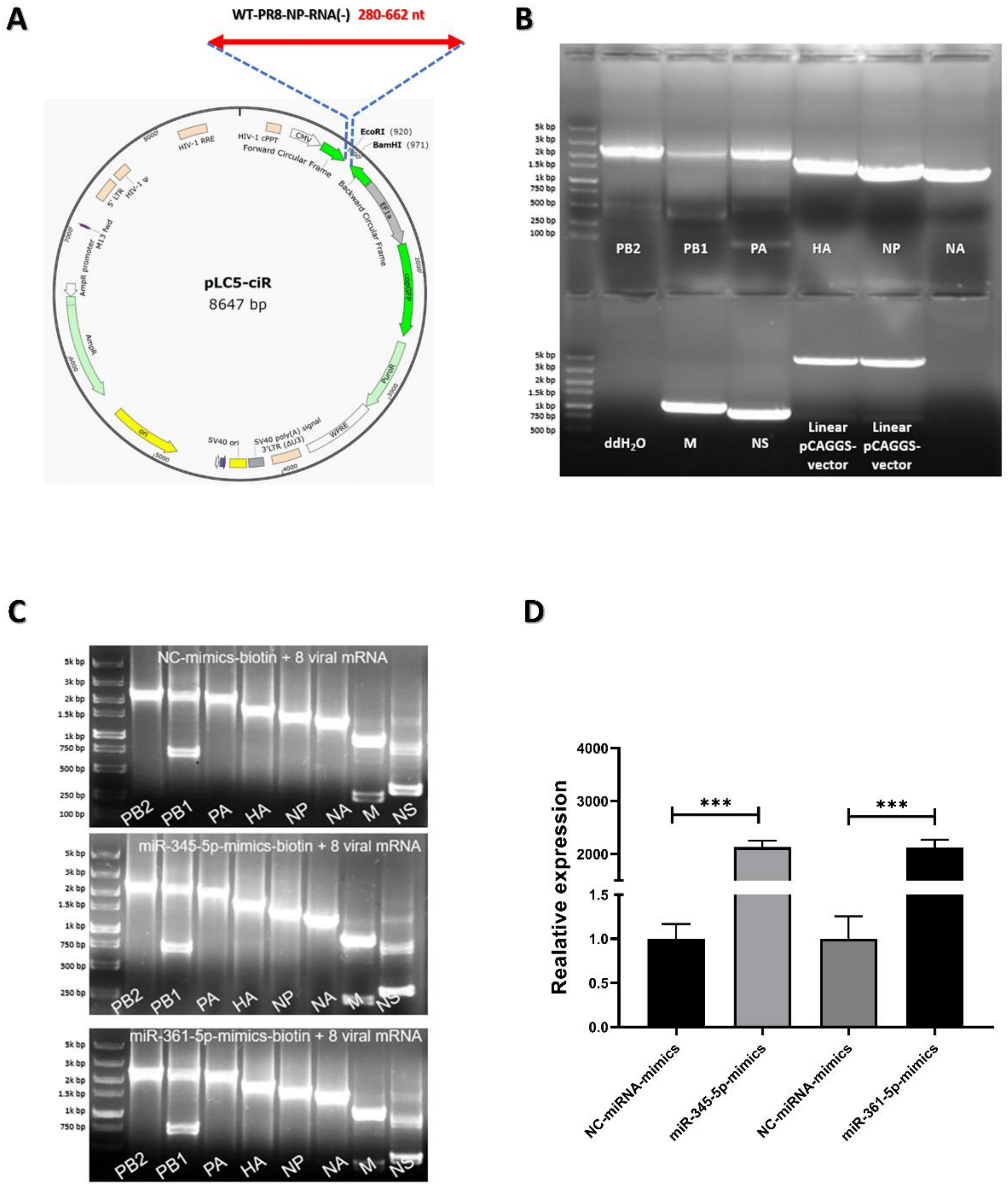


Figure S3:

A: CircNP37 expression plasmid.

B: Viral eight-gene product used to construct the pCAGGS expression plasmid.

C: Detection of viral mRNA expression by RT-PCR in miRNA-mimics cotransfection experiments with the pCAGGS expression vector expressing the viral 8 gene.

D: Detection of miR-345-5p and miR-361-5p overexpression efficiency in cells.

### KEY RESOURCES TABLE

| REAGENT or RESOURCE | SOURCE | IDENTIFIER |
|---|---|---|
| Antibodies | | |
| PB2 | GeneTex | Cat# GTX125926; RRID: AB_11162999 |
| NP | GeneTex | Cat# GTX125989; RRID: AB_11168364 |
| A-tubulin | Proteintech group, inc | Cat# 11224-1-AP-100UL; RRID: AB_2210206 |
| HRP-conjugated Goat Anti-Rabbit IgG | BBI | Cat# D110058-0001; |
| CircRNA Divergent Primers | | |
| circNP37-96-F | This paper | N/A |
| circNP37-96-R | This paper | N/A |
| circNP37-104-F | This paper | N/A |
| circNP37-104-R | This paper | N/A |
| circNP37-158-F | This paper | N/A |
| circNP37-158-R TGCAGTCAAAGGAGTTGGAACAATGG | This paper | N/A |
| circNP37-219-F AGTTCTCTCATCCACTTTCCG | This paper | N/A |
| circNP37-219-R TGTGCTCTCTGATGCAAGGTTC | This paper | N/A |
| circNP37-313-GSP ATTTGAATGATGCAACTTATCAG | This paper | N/A |
| circNP37-313-F TCATCATGTGAGTCAGACCAGCCGTTG | This paper | N/A |
| circNP37-313-R CAGGATGTGCTCTCTGATGCAAGGTTC | This paper | N/A |
| circNP37-367-F TGTTCCAACTCCTTTGACTGC | This paper | N/A |
| circNP37-367-R TCAGGATGATCAAACGTGGG | This paper | N/A |
| Viral Open Reading Frame Primers | | |
| PR8-PB2-F ATGGAAAGAATAAAAGAACTACGAAATCT | This paper | N/A |
| PR8-PB2-R CTAATTGATGGCCATCCGAATTC | This paper | N/A |
| PR8-PB1-F ATGGATGTCAATCCGACCTTAC | This paper | N/A |
| PR8-PB1-R CTATTTTTGCCGTCTGAGCTC | This paper | N/A |
| PR8-PA-F ATGGAAGATTTTGTGCGACAA | This paper | N/A |
| PR8-PA-R CTAACTCAATGCATGTGTAAGGAAG | This paper | N/A |
| PR8-HA-F ATGAAGGCAAACCTACTGGTC | This paper | N/A |
| PR8-HA-R TCAGATGCATATTCTGCACTGC | This paper | N/A |
| PR8-NP-F ATGGCGTCTCAAGGCACCAAA | This paper | N/A |
| PR8-NP-R TTAATTGTCGTACTCCTCTGCATT | This paper | N/A |
| PR8-NA-F ATGAATCCAAATCAGAAAATAATAACCAT | This paper | N/A |
| PR8-NA-R CTACTTGTCAATGCTGAATGGC | This paper | N/A |
| PR8-M-F ATGAGTCTTCTAACCGAGGTCG | This paper | N/A |
| PR8-M-R TTACTCCAGCTCTATGCTGACAA | This paper | N/A |
| PR8-NS-F ATGGATCCAAACACTGTGTCA | This paper | N/A |
| PR8-NS-R CTAAATAAGCTGAAACGAGAAAGTTCT | This paper | N/A |
| miRNA Primers | | |
| U6-miRNA-F CTCGCTTCGGCAGCACA | This paper | N/A |
| U6-miRNA-R AACGCTTCACGAATTTGCGT | This paper | N/A |
| hsa-miR-361-5p-GSP GTCGTATCCAGTGCAGGGTCCGAGGTATTCGCACTGGATACGACGTACCC | This paper | N/A |
| hsa-miR-361-5p-F GCGCGTTATCAGAATCTCCAG | This paper | N/A |
| hsa-miR-345-5p-GSP GTCGTATCCAGTGCAGGGTCCGAGGTATTCGCACTGGATACGACGAGCCC | This paper | N/A |
| hsa-miR-345-5p-F GCGGCTGACTCCTAGTCCA | This paper | N/A |

(*Continued on next page*)

| Continued | | |
|---|---|---|
| REAGENT or RESOURCE | SOURCE | IDENTIFIER |
| hsa-miR-769-3p-GSP GTCGTATCCAGTGCAGGGTCCGAGGTATTCGCACTGGATACGACAACCAA | This paper | N/A |
| hsa-miR-769-3p-F GCTGGGATCTCCGGGGTC | This paper | N/A |
| miRNA-R AGTGCAGGGTCCGAGGTATT | This paper | N/A |
| Quantitative Real-time PCR Primers | | |
| Influenza A virus U-12_GSP (A) AGCGAAAGCAGG | This paper | N/A |
| Influenza A virus U-12_GSP (B) AGCAAAAGCAGG | This paper | N/A |
| PB2-qPCR-143-F CCAAGTCAAAATACGTCGGAG | This paper | N/A |
| PB2-qPCR-143-R TCGTTAGTTGCGATTCCGAT | This paper | N/A |
| PB1-qPCR-186-F TAATTAGAGCATTGACCCTGAACACA | This paper | N/A |
| PB1-qPCR-186-R CTTTCTTCTCATTGCCTCCAACTG | This paper | N/A |
| PA-qPCR-100-F GCTTCTTATCGTTCAAGCTCTT | This paper | N/A |
| PA-qPCR-100-R GGGATCATTAATTAGGCACTCC | This paper | N/A |
| HA-qPCR-102-F GGCCCAACCACAACACAAAC | This paper | N/A |
| HA-qPCR-102-R AGCCCTCCTTCTCCGTCAGC | This paper | N/A |
| NP-qPCR-181-F CACTTTCCGTTTACTCTCCTGTATATAGG | This paper | N/A |
| NP-qPCR-181-R GCACCGAACTTAAACTCAGTGATTATGAG | This paper | N/A |
| NA-qPCR-144-F CGGCCATGGGTGTCTTTC | This paper | N/A |
| NA-qPCR-144-R TCCCTTTACTCCGTTTGCTCCATC | This paper | N/A |
| M-qPCR-95-F AGGATGGGGGCTGTGACC | This paper | N/A |
| M-qPCR-95-R ATTTGCCTATGAGACCGATGCT | This paper | N/A |
| NS-qPCR-155-F GATAGTGGAGCGGATTCTGA | This paper | N/A |
| NS-qPCR-155-R GAGGGCCTGCCACTTTCT | This paper | N/A |
| U6-70-F ATTGGAACGATACAGAGAAGATT | This paper | N/A |
| U6-70-R GGAACGCTTCACGAATTTG | This paper | N/A |
| Human GAPDH endogenous reference genes primers | BBI | B661104-0001 |
| Human ACTB endogenous reference genes primers | BBI | B661102-0001 |
| Mouse GAPDH endogenous reference genes primers | BBI | B661304-0001 |
| Molecular Cloning Primers | | |
| 5’ bss-mutant-F CTGGGATGTTCTTCCAaaTATTTATTTCTCCTTTCGTCAAAAGC | This paper | N/A |
| 5’ bss-mutant-R ttTGGAAGAACATCCCAGTGCG | This paper | N/A |
| 3’ bss-mutant-F ACCCCTCCAaAAGTTCCGATCATTGATCCCAC | This paper | N/A |
| 3’ bss-mutant-R GGAACTTtTGGAGGGGTGAGAATGGACG | This paper | N/A |
| PB2 homologous recombination 2393F TGTCTCATCATTTTGGCAAAGAATTCagcaaaagcaggtcaattatattcaat | This paper | N/A |
| PB2 homologous recombination 2393R GAGGGAAAAAGATCTGCTAGCTCGAGagtagaaacaaggtcgtttttaaactattc | This paper | N/A |
| PB1 homologous recombination 2393F TGTCTCATCATTTTGGCAAAGAATTCagcgaaagcaggcaaacc | This paper | N/A |
| PB1 homologous recombination 2393R GAGGGAAAAAGATCTGCTAGCTCGAGagtagaaacaaggcattttttcatgaagga | This paper | N/A |
| PA homologous recombination 2285F GTCTCATCATTTTGGCAAAGAATTCagcgaaagcaggtactgatccaaa | This paper | N/A |

(*Continued on next page*)

| Continued | | |
|---|---|---|
| REAGENT or RESOURCE | SOURCE | IDENTIFIER |
| PA homologous recombination 2285R GAGGGAAAAAGATCTGCTAGCTCGAGagtagaaacaaggtactttttggacagtatgg | This paper | N/A |
| HA homologous recombination 1830F TGTCTCATCATTTTGGCAAAGAATTCagcaaaagcaggggaaaataaaaacaacc | This paper | N/A |
| HA homologous recombination 1830R GAGGGAAAAAGATCTGCTAGCTCGAGagtagaaacaagggtgtttttcctcatat | This paper | N/A |
| NP homologous recombination 1617F TGTCTCATCATTTTGGCAAAGAATTCagcaaaagcagggtagataatcactcac | This paper | N/A |
| NP homologous recombination 1617R GAGGGAAAAAGATCTGCTAGCTCGAGagtagaaacaagggtatttttctttaat | This paper | N/A |
| NA homologous recombination 1465F TGTCTCATCATTTTGGCAAAGAATTCagcgaaagcaggagtttaaaatgaat | This paper | N/A |
| NA homologous recombination 1465R GAGGGAAAAAGATCTGCTAGCTCGAGagtagaaacaaggagtttttgaacaga | This paper | N/A |
| M homologous recombination 1079F TGTCTCATCATTTTGGCAAAGAATTCagcaaaagcaggtagatattgaaag | This paper | N/A |
| M homologous recombination 1079R GAGGGAAAAAGATCTGCTAGCTCGAGagtagaaacaaggtagtttttactcc | This paper | N/A |
| NS homologous recombination 942F TGTCTCATCATTTTGGCAAAGAATTCagcaaaagcagggtgacaaaaacat | This paper | N/A |
| NS homologous recombination 942R GAGGGAAAAAGATCTGCTAGCTCGAGagtagaaacaagggtgtttttattatt | This paper | N/A |
| circNP37-miR-345-5p-seed region-Mut-F CATGTctcagtcACCAGCCGTTGCATCGTCAC | This paper | N/A |
| circNP37-miR-345-5p-seed region-Mut-R CTGGTgactgagACATGATGATCTGGCATTCCAA | This paper | N/A |
| circNP37-miR-361-5p-seed region-Mut-F TGTCCagactattGTTGCATCATTCAAATTGGAATGC | This paper | N/A |
| circNP37-miR-361-5p-seed region-Mut-R GCAACaatagtctGGACAAGAGCTCTTGTTCGCA | This paper | N/A |
| Probes | | |
| circNP37_5' biotin_probe_1 ACTTCTGGAAGAACATCCCA | RiboBio | N/A |
| circNP37_5' biotin_probe_2 GAACTTCTGGAAGAACATCC | RiboBio | N/A |
| circNP37_5' biotin_probe_3 CGGAACTTCTGGAAGAACAT | RiboBio | N/A |
| circNP37_5' biotin_probe_Scramble ChIRP TAAGTGCTCGTAAGCAACCA | RiboBio | N/A |
| PB2-mRNA_5' biotin_probe_1 TGCCATCATCCATTTCATCC | RiboBio | N/A |
| PB2-mRNA_5' biotin_probe_2 TTTGTCCTTGCTCATTTCTC | RiboBio | N/A |
| miRNA mimics/miRNA inhibitor | | |
| miR-361-5p-mimics | Beijing Tsingke Biotech | N/A |
| miR-361-5p-inhibitor | Beijing Tsingke Biotech | N/A |
| miR-345-5p-mimics | Beijing Tsingke Biotech | N/A |
| miR-345-5p-inhibitor | Beijing Tsingke Biotech | N/A |
| NC-mimics | Beijing Tsingke Biotech | N/A |
| NC-inhibitor | Beijing Tsingke Biotech | N/A |
| Biotin-miR-361-5p-mimics | Beijing Tsingke Biotech | N/A |
| Biotin-miR-345-5p-mimics | Beijing Tsingke Biotech | N/A |
| Biotin-NC-mimics | Beijing Tsingke Biotech | N/A |
| Silencer | | |
| Silencer_1 GTTCTTCCAGAAGTTCCGAT | RiboBio | N/A |

(*Continued on next page*)

| Continued | | |
|---|---|---|
| REAGENT or RESOURCE | SOURCE | IDENTIFIER |
| Silencer_2 CTTCCAGAAGTTCCGATCAT | RiboBio | N/A |
| Silencer_3 GGATGTTCTTCCAGAAGTTC | RiboBio | N/A |
| Silencer_4 GGATGTTCTTCCAGAAGTT | RiboBio | N/A |
| Silencer_5 GTTCTTCCAGAAGTTCCGA | RiboBio | N/A |
| Silencer_6 CTTCCAGAAGTTCCGATCA | RiboBio | N/A |
| Cell Culture | | |
| Opti-MEM™ Medium | Gibco™ | Cat# 31985070 |
| Penicillin - Streptomycin (5,000 U/mL) | Gibco™ | Cat# 15070063 |
| PBS | BBI | Cat# E607008-0500 |
| Dulbecco's modified Eagle's medium (DMEM) | Gibco™ | Cat# C11995500BT |
| Fetal bovine serum | Gibco™ | Cat# 10099141C |
| 1-Toluenesulfonamide-2-phenylethyl chloromethyl ketone (TPCK)-treated trypsin | Sigma-Aldrich | Cat# T1426 |
| Bovine serum albumin | BBI | Cat# A600332-0100 |
| Critical Commercial Assays and kits | | |
| Lipofectamine™ 3000 Reagent | Invitrogen™ | Cat# L3000015 |
| Phanta Max Super-Fidelity DNA Polymerase | Vazyme | Cat# P505-d1 |
| EcoRI | NEB | Cat# R0101V |
| XhoI | NEB | Cat# R0101V |
| FastPure Gel DNA Extraction Mini Kit | Vazyme | Cat# DC301-01 |
| ClonExpress II One Step Cloning Kit | Vazyme | Cat# C112-02 |
| DH5α competent cell | Vazyme | Cat# C502-03 |
| Ampicillin | Sangon Biotech | Cat# B541011-0001 |
| Hieff Trans® in vitro siRNA/miRNA Transfection Reagent | YEASEN | Cat# 40806ES01 |
| EndoFree Mini Plasmid Kit II | TIANGEN | Cat# DP118 |
| Polyethylenimine Linear (PEI) MW25000 | YEASEN | Cat# 40815ES03 |
| Mut Express II Fast Mutagenesis Kit V2 | Vazyme | Cat# C214-02 |
| RNA isolater Total RNA Extraction Reagent | Vazyme | Cat# R401-01 |
| RNase-Free dd$H_2O$ | Sangon Biotech | Cat# B541018-0010 |
| HiScript® III 1st Strand cDNA Synthesis Kit (+gDNA wiper) | Vazyme | Cat# R312-01 |
| 2×TSINGKE® Master Mix (Blue) | TsingkeBiotechnology | Cat# TSE004 |
| Taq Pro Universal SYBR qPCR Master Mix | Vazyme | Cat# Q712 |
| RNase R | GENESEED | Cat# R0301 |
| 10×Poly-L-Lysine | Solarbio | Cat# P2100-10mL |
| Hieff Trans® Liposomal Transfection Reagent | YEASEN | Cat# 40802ES03 |
| ExFect Transfection Reagent | Vazyme | Cat# T101-01 |
| Cytoplasmic & Nuclear RNA Purification Kit | NORGEN | Cat# 21000 |
| BersinBio™ RNA Antisense Purification (RNA pull-down) Kit | BersinBioTM | Cat# Bes5103-3 |
| Phenol-chloroform-isoamyl alcohol mixture (25:24:1) | Solarbio | Cat# P1011-100mL |
| miRNA pulldown Kit | BersinBioTM | Cat# Bes5108 |
| Dual Luciferase Reporter Assay Kit | Vazyme | Cat# DL101 |
| RIPA | Beyotime | Cat# P0013B |
| PMSF | Beyotime | Cat# ST506 |
| 5 × Loading Buffer | Beyotime | Cat# P0015L |
| One-Step PAGE Gel Fast Preparation Kit (10%) | Vazyme | Cat# E303 |
| Western Rapid Transfer Buffer (10X) | Beyotime | Cat# P0572 |

(*Continued on next page*)

| Continued | | |
|---|---|---|
| REAGENT or RESOURCE | SOURCE | IDENTIFIER |
| 10 × TBS | Beyotime | Cat# ST661 |
| skimmed milk powder | Beijing Dingguo | Cat# DH220 |
| 180 kDa Prestained Protein Marker | Vazyme | Cat# MP102-01 |
| 100 bp DNA Ladder | Vazyme | Cat# MD104-01 |
| DL2000 Plus DNA Marker | Vazyme | Cat# MD101-01 |
| DL5000 DNA Marker | Vazyme | Cat# MD102-01 |
| DL15000 DNA Marker | Vazyme | Cat# MD103-01 |
| Western blotting, dot/slot blotting | Millipore | Cat# WBKLS0100 |
| Avertin | This paper | N/A |
| 4% Paraformaldehyde Fix Solution | Beyotime | Cat# P0099-100mL |
| Experimental modles:Cells | | |
| A549 cells (Non-small cell lung cancer cell lines) | Laboratory preservation | N/A |
| HEK 293T cells (human embryonic kidney 293T cells) | Laboratory preservation | N/A |
| MDCK cells (madin-darby canine kidney) | Laboratory preservation | N/A |
| Experimental modles:Organisms | | |
| Mouse: BALB/c | Beijing Weitong Lihua Laboratory Animal Technology Co. | N/A |
| Plasmid | | |
| Influenza A virus (PR8) Reverse Genetic System | Laboratory preservation | N/A |
| 5'+3' bss-pHW2000-NP | This paper | N/A |
| pLC5-circNP37 | GENESEED | N/A |
| pLC5 | GENESEED | N/A |
| pLC5-circNP37-miR-361-5p-Mut-Site | This paper | N/A |
| pmiRGLO-circNP37 | GUANGZHOU IGE BIOTECHNOLOGY LTD | N/A |
| pmiRGLO-circNP37-miR-361-5p-Mut-Site | This paper | N/A |
| pmiRGLO-circNP37-miR-345-5p-Mut-Site | This paper | N/A |
| pCAGGS-PB2 | This paper | N/A |
| pCAGGS-PB1 | This paper | N/A |
| pCAGGS-PA | This paper | N/A |
| pCAGGS-HA | This paper | N/A |
| pCAGGS-NP | This paper | N/A |
| pCAGGS-NA | This paper | N/A |
| pCAGGS-M | This paper | N/A |
| pCAGGS-NS | This paper | N/A |
| Soferware | | |
| GraphPad Prism 8 | GraphPad Software | https://www.graphpad-prism.cn/ |
| ImageJ | NIH, USA | https://imagej.nih.gov/ij/ |
| Snapgene Viewer | Snapgene | https://www snapgene com/ |
| miRbase | Ana Kozomara et al., 2019 | https://www.mirbase.org/ |
| miRanda | A.J. Enright et al., 2003 | http://www.bioinformatics.com.cn/local_miranda_miRNA_target_prediction_120 |
| miRDB | Yuhao Chen and Xiaowei Wang, 2020 | https://mirdb.org/mirdb/index.html |
| Venn diagram | Philippe Bardou et al., 2014 | http://www.bioinformatics.com.cn/static/others/jvenn/example.html |
| RNAhybird | Jan Krüger and Marc Rehmsmeier, 2006 | https://bibiserv.cebitec.uni-bielefeld.de/rnahybrid |
| miRmine | Bharat Panwar et al.,2017 | https://guanfiles.dcmb.med.umich.edu/mirmine/index.html |

#### LEAD CONTACT AND MATERIALS AVAILABILITY

#### EXPERIMENTAL MODEL AND SUBJECT DETAILS

##### Cell lines

A549 cells, 293T cells, and MDCK cells were maintained in this laboratory. These three cell lines were cultured in Dulbecco's modified Eagle's medium (DMEM) (Gibco™) containing 10% fetal bovine serum (Gibco™), 2% penicillin-streptomycin (5,000 U/mL) (Gibco™). Cells were cultured at 37 °C in 5% $CO_2$. Passaging cultures were performed when cells reached 80%-90% confluence of growth.

##### Mice

The experiments were performed in accordance with the Chinese national guidelines for the care of laboratory animals. 5-7 week-old SPF-grade female BALB/c mice were purchased from Beijing Viton Lever Laboratory Animal Technology Ltd (Beijing, China) and housed in an animal biosafety level 2 facility at the Wuhan Institute of Virology, Chinese Academy of Sciences (Wuhan, China). In vivo experiments were approved by the Animal Welfare and Ethical Review Committee of Wuhan Institute of Virology, Chinese Academy of Sciences (WIVA39202301).

##### Viruses

Influenza A virus H1N1 (A/Puerto Rico/8/34, H1N1 [PR8]) was rescued by the viral reverse genetics system, and the viral genome sequence information is shown in the table 1. The virus reverse genetic system is based on the pHW2000 plasmid, which is maintained in our laboratory. Transfection of pHW2000 with viral genes in 293T cells using Lipofectamine™ 3000 Reagent (Invitrogen™).The IAV rescue process, A: 500 ng of each plasmid (4 μg total), mixed with 8 μL of P3000™ Reagent in 125 μL of Opti-MEM™ Medium (Gibco™); B: 7.5

μL of Lipofectamine™ 3000 Reagent (Invitrogen™) mixed with 125 μL of Opti-MEM™ Medium; Mix A and B thoroughly for 15 min at room temperature, add to six-well plate and supplement with 500 μL Opti-MEM™ Medium, incubate for 5-6 h at 37 °C, then supplement with 2 mL Opti-MEM™ Medium and continue incubation at 37 °C. After 48 hours, the cell supernatant was collected, filtered through a 0.22 μm filter, and then injected into 9-11 day-old SPF chicken embryos (Seth Poultry Technology Co., Ltd., Jinan, China). The virus-containing chicken embryo allantoic fluid was collected after 72 hours, filtered through a 0.22 μm filter again, and then dispensed, and frozen at -80 °C.

**Table 1 Genome sequences of virus reverse genetics system**

| | Viral gene | GenBank_ACCESSION |
|---|---|---|
| 1 | PB2 | MH785018 |
| 2 | PB1 | MH785017 |
| 3 | PA | MH785016 |
| 4 | HA | MH785011 |
| 5 | NP | MH785014 |
| 6 | NA | MH785013 |
| 7 | M | MH785012 |
| 8 | NS | MH785015 |

### METHOD DETAILS

**Viral hemagglutination titer assay**

A V-shaped 96-well hemagglutination plate was used for the assay. Add 90 μL of PBS (BBI) to the first well and 50 μL of PBS to each of the second to twelfth wells. 10 μL of the sample to be tested was added to the first well, gently mixed, then take 50 μL was added to the second well, 50 μL was mixed and added to the third well, and so on, and after dilution to the tenth well, the diluted sample was discarded. The experiment's final two wells serve as negative controls. After 50 μL of 1% chicken erythrocyte suspension (prepared in our laboratory) was added dropwise to each well, mixed gently, and left at room temperature for 20-25 min, the hemagglutination plate was stood up vertically, turned over to the back side, and the viral hemagglutination titer was recorded.

**Plasmid construction**

pCAGGS expression vector and pHW2000 Virus Reverse Genetics System is maintained in our laboratory. pLC5-circNP37 plasmid was purchased from Geneseed Biotechnology Co., LTD (Guangzhou, China); pmiRGLO-circNP37 was purchased from GUANGZHOU IGE BIOTECHNOLOGY LTD (Guangzhou, China).

The pCAGGS vector containing the viral 8 gene mRNA was constructed for this experiment, and the full-length viral positive-strand RNA amplification primers with homologous arms were designed based on the flanking sequence of the pCAGGS digestion site. The viral positive-stranded sequence fragment (PB2, PB1, PA, HA, NP, NA, NP, M, NS) was amplified using cDNA produced by reverse transcription of RNA from Chicken embryo allantoic fluid containing the virus. The pCAGGS expression vector was double digested (37 °C, > 12 h) using EcoRI (NEB) as well as XhoI (NEB), and the linear vector was purified using FastPure Gel DNA Extraction Mini Kit (Vazyme) gel recovery. The viral positive strand sequence with the homologous arm was cloned into the pCAGGS plasmid by homologous recombination using the ClonExpress II One Step Cloning Kit (Vazyme) according to the instructions. Recombination products were transfected into DH5α competent cells (Vazyme). Plasmid extraction was performed using the EndoFree Mini Plasmid Kit II (TIANGEN). The extracted plasmids were sent to TsingkeBiotechnologyCo., Ltd. for sequencing and validation. The pCAGGS expression vector containing viral 8 gene mRNA was transfected in 293T cells with 625 ng of each plasmid using PEI25000 (YEASEN) 10 μL, and total RNA was extracted from the cells after 24 h. The viral ORF sequence was amplified using RT-PCR to detect viral 8 gene mRNA transcription. The plasmid was placed into experimental use when it was confirmed that it was able to properly express viral 8 gene mRNA.

**Point mutation**

Using Mut Express II Fast Mutagenesis Kit V2 (Vazyme), the 5'+3'bss mutated pHW2000-NP plasmid, miR-345-5p, miR-361-5p seed region altered pmiRGLO-circNP37 plasmid, and pLC5-circNP37-Mut (altered miR-361-5p seed region) were constructed. In brief, PCR amplification using plasmid as templates, producing in a linearized fragment that is recirculated into an entire plasmid via homologous recombination. The recombinant products were transfected into DH5α competent cells. Plasmid extraction was followed by sequencing to verify successful cloning.

**RNA extraction**

RNA isolater Total RNA Extraction Reagent (Vazyme) was used for RNA extraction. The supernatant from the six-well plate was removed, and 1 mL of RNA Extraction Reagent was added. Ttransfer the lysed cells to an EP tube, and let stand for 5 min at 4 ℃. Add chloroform to the lysate (1 mL RNA Extraction Reagent to 0.2 mL chloroform), and vortex vigorously for 15 s. Centrifuge at 12,000 rpm for 15 min at 4 ℃. Transfer the aqueous phase (colorless supernatant) to a new EP tube. To the colorless supernatant, add an equal volume of pre-cooled isopropanol, mix well, and store at 4 ℃ for 10 minutes. Centrifuge at 12,000 rpm for 10 min at 4 ℃ and discard the supernatant. Add 75% ethanol (RNase-Free) to each EP tube and wash the white precipitate. Remove the supernatant and centrifuge at 12,000 rpm for 3 min at 4 ℃. Allow EP tube caps to dry at room temperature for 2-3 minutes before adding 30-100 μL of RNase-free $ddH_2O$ (Sangon Biotech).

**Reverse Transcript-Polymerase chain reaction (RT-PCR) and Real-time Quantitative PCR**

In the reverse transcription assays, random primers, Oligo (dT), and Gene Specific Primer (GSP) were used according to the experimental requires. In this experiment, the reverse transcription of circRNA was performed by GSP. The HiScript® III 1st Strand cDNA Synthesis Kit (+gDNA wiper) (Vazyme) was used for the reverse transcription reaction.

For circNP37 BSJ PCR amplification, the reaction system was prepared using 2×TSINGKE® Master Mix (Blue) (Tsingke Biotechnology) and the Touch Down PCR procedure was used for Amplification (annealing temperature: 62-56℃, 12 cycles, -0.5℃/cycles; 56℃, 23 cycles). The PCR products were detected by electrophoresis. Voltage and current: 160 V, 230 mA, 30 min. PCR products were collected and sent to Tsingke Biotech for sequencing. For circNP37 amplification products, the fragments were cloned into plasmid vectors and sequenced to determine the exact nucleotide makeup.

In the qPCR experiment, the reaction system was prepared using Taq Pro Universal SYBR qPCR Master Mix (Vazyme). The relative quantification, $2^{-\Delta\Delta}$ Ct method, was used for the calculation of the experiments.

For viral copy number calculation, absolute quantitative PCR was used for the assay. The pHW2000-NP, pHW-2000-M and pCAGGS-PB2 plasmids were used as standards in the experiments, respectively. Following the determination of plasmid concentrations, the number of molecular copies per microliter in the undiluted samples was estimated separately, and seven dilutions of each plasmid were done in a tenfold gradient dilution, yielding standards for NP, M, and PB2, at $10^{-1}$-$10^{-7}$ dilutions. The qPCR was performed separately with the standard as the template and the sample to be tested, and the copy number of molecules contained in the sample to be tested was calculated using the standard curve.

**RNase R digestion assay**

The reaction system was set up in accordance with the instructions (Geneseed). 5 μg of RNA was put into the reaction system, and RNase R (Geneseed) was added 10 U, digested at 37 °C for 10 minutes, and RNase R inactivated at 70 °C for 10 minutes. Subsequent reverse transcription, qPCR assays were performed as described in the previous section.

**Virus Infections**

Configuration of virus adsorption solution: DMEM with 1 μg/mL L-1-toluenesulfonamide-2-phenylethyl chloromethyl ketone (TPCK)-treated trypsin (Sigma-Aldrich) and 0.3% Bovine serum albumin (BBI). Prepare for each experiment the day before infection by adding equal amounts of diluted cell suspension to each well (6-well plate). For viral infection, cells are are cultured to approximately 70-80% confluence. All cells in the cell counting wells are collected and counted. MOI= PFU/cells, TCID50×0.7≈ PFU, converted in this way, and virus dilution was performed according to the amount of virus required for the experiment. At the time of infection, the cell supernatant in the 6-well plate was discarded, PBS was added and gently washed twice, diluted virus was added, and one well of the six-well plate was used for virus adsorption in a volume of 600 μL. The culture dish was gently shaken every 20 minutes while the diluted virus was injected for around 2 hours, then, the cell supernatant in the wells was removed, PBS was gently washed once, and 2 mL of virus adsorption solution was added.

**Virus titer determination**

Median tissue culture infective dose (TCID50) assay: MDCK cells were cultured in 96-well plate the day before infection. After the cells in the 96-well plate have grown to a density of about 80-90%, the TCID50 infection assay is performed. The virus to be tested was subjected to a ten-fold gradient dilution. Discard the 96-well plate supernatant, wash gently twice with PBS, and then remove the PBS. The 100 μL of diluted virus was added to each well. The 96-well plate was placed in the culture incubator for around 1-2 hours after the virus was injected. The 96-well plate was gently shaken every 20 minutes during this time to promote virus entry into the cells. At the end of the infection, the cell supernatant in each well was removed, the PBS washed once, and the PBS was discarded. Adding 100 μL of viral adsorbent to the 96-well plate, the culture was continued for 72 hours, with cytopathic effect being monitored every 24 hours. After 72 h, the virus titer was calculated by the Reed-Muench method.

**Expression of circNP37 during Mut-PR8 infection**

293T cells were cultured in 6-well plate the day before transfection. 6-well plate. Lipofectamine™ 3000 Reagent (Invitrogen™) was used to transfect the circNP37 plasmid (5 μg) into the cells, in accordance with the instruction. 12 h after plasmid transfection, viral infection was performed as described in the previous part of the viral infection experiment. After 24 h of infection, total cellular RNA was extracted for reverse transcription and qPCR assay, and cell supernatants were collected for virus titer assay.

**WT-PR8 knockdown of circNP37 during infection**

293T cells were cultured in 6-well plate the day before transfection. Silencer transfection was performed using Hieff Trans® in vitro siRNA/miRNA Transfection Reagent (YEASEN). Mix 7.5 μL of silencer (20 μM), 200 μL of Opti-MEM™ Medium (Gibco™), and 5 μL of Hieff Trans® in vitro siRNA/miRNA Transfection Reagent and incubate at room temperature for 10 min. Cell supernatants were removed, washed once with PBS (BBI), and DMEM (Gibco™) was added to bring the final Silencer transfection concentration to 150 nM. After 12 h of transfection, viral infection was performed as described previously. After 36 h of virus infection, total cellular RNA was extracted for qPCR detection, and cell supernatants were collected for virus titer assay.

**miR-361-5p-mimics/inhibitor transfection followed by plasmid transfection or viral infection**

The Hieff Trans® Liposomal Transfection Reagent (YEASEN), miRNA-mimics, and Opti-MEM temperatures were balanced to room temperature before miRNA-mimics transfection was performed out on 293T cells in 6-well plate. Hieff transfection reagent and 120 mL of Opti-MEM were mixed gently and let to stand for 5 min at room temperature. Place 10 μL/15 μL of miR-361-5p-mimics/miR-361-5p-inhibitor at a concentration of 20 μM in the transfection system and let stand at room temperature for 10 min. During this period, discard the cell supernatant, add PBS and wash gently once, discard the PBS, add serum-free and antibiotic-free DMEM and transfection complex to reach a final concentration of 150 nM for miRNA-mimics and 200 nM for miRNA-inhibitor. The final concentration of miRNA-mimics in the six-well plate reaches 150 nM and the final concentration of miRNA-inhibitor reaches 200 nM. Plasmid transfection was performed after 12 h. The plasmids were gently mixed according to Opti-MEM 400 μL, ExFect Transfection Reagent (Vazyme) 15 μL and 5 μg of plasmid and left for 15 min at room temperature. The cell supernatant was removed, 1.6 mL of DMEM was added, and the plasmid complex was gently added dropwise to a six-well plate and placed in 37 °C incubation. Transfected for 24-48h for assay. Alternatively, the miRNA-mimics/inhibitor was transfected for 8 h, WT-PR8/Mut-PR8 MOI=0.01 was infected, and the experimental samples were collected at 48 hpi for detection.

**Nuclear/Cytoplasmic Fractionation**

Purification of nuclear and cytoplasmic RNA was performed using Cytoplasmic & Nuclear RNA Purification Kit (NORGEN). A549 cells were cultured in 6-well plate the day before the WT-PR8 infection. The cells were grown to ~80% confluence and viral infection was performed. Experiments were performed at MOI = 1. At 24 hpi, the cell supernatant was discarded and the cells were gently washed once with ice-cold PBS. After cell lysis, the cytoplasmic and nuclear fractions were separated by centrifugation, and after purification of the two RNA fractions separately by sponge column. qPCR was performed as described above.

**RNA Antisense Purification**

In experiments, RNA is pull-down using a biotin-labeled probe; if other nucleic acids or proteins that interact with the RNA are present, they can also be pull-down simultaneously. To lessen the chance of off-targeting, three oligonucleotide probes (5' biotin-labeled) with different sequences were designed for circRNA BSJ in this experiment. Three distinct probes were employed simultaneously for circRNA pull-down. For PB2-mRNA, two probes with different sequences were used simultaneously for pull-down in one experiment. The scrambled oligonucleotide probe pull-down group was used as the negative control in the experiment. BersinBio™ RNA Antisense Purification (RNA pull-down) Kit (BersinBio™) was used for this experiment. Transfection experiments were performed with Polyethylenimine Linear (PEI) MW25000 (YEASEN). Mix together 4 mL of Opti-MEM™ Medium (Gibco™), 150 μL of PEI 25000 (YEASEN), and 50 μg of pLC5-circNP37 and allow to stand at room temperature for 15 minutes. 293T cell supernatant was removed and gently washed once with PBS, PBS was removed, serum-free and antibiotic-free DMEM medium (Gibco™) was added, and the transfection. After 48 h RAP experiments were performed.

In the RAP experiment, cells were first collected from the culture dish, 10 mL of ice-cold PBS was added, and the cells were washed. Add 40 mL of PBS containing a final concentration of 1% formaldehyde to the cells and mix for 10 min on a vertical mixer. Add 4 mL of 1.375 M Glycine and mix for 5 min on a vertical mixer. Centrifuge at 1500 rpm at 4 °C for 5 min to remove the supernatant, and then add 10 mL of ice-cold PBS to wash the cells twice under the same conditions as before. Mix 1.8 mL of Lysis buffer, 18 μL of protease inhibitor and 9 μL of RNase Inhibitor add to the cell precipitate, vortex and mix well, and place on ice for 10 min for lysis. Take 9 μL DNase salt stock, 20 μL DNase (20 U), mix well and add to the cell lysate and mix thoroughly. Add equal volumes of 2×Hybridization Buffer to the probe pull-down group and negative control group, respectively, mix well and put on ice for 10 min. For hybridization of magnetic beads and probes, denature the lysate samples at 65 °C for 10 min, add circNP37 probe and scrambled control probe, respectively, and hybridize at 37 °C for 3 h. Place the hybridized sample in the prepared magnetic beads and mix on a vertical mixer for 30 min. After removing the supernatant, add 500 μL of elution buffer, elute at 37 °C for 5 min, remove the supernatant, and repeat this step three times. After removal of the supernatant add 50 μL of elution buffer, process at 95 °C for 2 min, place the sample on a magnetic separation device, transfer the supernatant to a clean EP tube and repeat the step once. Add 25 μL 5×Proteinase K buffer and 2 μL proteinase K to the sample, and digest at 55 °C for 1 h. An equal volume of phenol-chloroform-isoamyl alcohol mixture (25:24:1) (Solarbio) was added to the digested samples and mixed upside down for 15 s. The samples were centrifuged at 13,000×g for 10 min, and the aqueous phase of the supernatant was transferred to a new RNase-free tube. Add 5 μL of NaCl, 1 μL of Glycogen, 312.5 μL of anhydrous ethanol, mix upside down, and precipitate at -80 °C for 12 h.

Centrifuge the RNA sample at 16000×g for 30 min after 12 h precipitation, discard the supernatant and dry at room temperature, add 30 μL of RNase-free $ddH_2O$ (Sangon Biotech). The reverse transcription and qPCR experiments were performed as described previously, and the differential changes in expression of circNP37, GAPDH, U6 snRNA and candidate miRNAs were quantified. PB2-mRNA RAP experiments protocol were performed as in the circNP37 RAP experiments protocol.

**miRNA pull down**

miRNA pull down experiments were performed using the miRNA pulldown Kit (BersinBio™). The experimental plan was to overexpress viral mRNAs in cells while transfection miRNA-mimics with 5' biotin tags. Prior to transfection experiments, 293T cells were cultured in 15 $cm^2$ cell culture dishes, (2 15 $cm^2$ cell culture dishes were prepared in a set of miRNA pull-down experiments). When the cells grew to ~80% confluence, miRNA-mimics transfection was performed. The miRNA transfection was performed using Hieff Trans® Liposomal Transfection Reagent (YEASEN). Mix 1850 μL of Opti-MEM (Gibco™) with 107.919 μL of Hieff Trans™ Liposomal Transfection Reagents and let stand at room temperature for 5 min. Add 154.17 μL of biotin-labeled miRNA-mimics (20 μM) to the above mixture and let stand at room temperature for 10 min. The cell supernatant was replaced with 17887 μL of DMEM, and the miRNA transfection mix were added dropwise to the dishes and mixed gently. Transfection of plasmids PB2, PB1, PA, HA, NP, NA, M, and NS was performed 6 h after miRNA transfection at a total transfection dose of 37.5 μg, with the same amount of transfection for each plasmid. The plasmids were mixed with 6 mL of Opti-MEM (Gibco™) and 225 μL of PEI 25000 was added. Cells were first treated with 0.25% Trypsin-EDTA (Gibco™) for 3 min and then collected and washed twice with PBS. After cell precipitation was collected, the supernatant was removed as much as possible and stored frozen at -80 °C.

To perform bead closure, 40 μL of streptavidin beads were washed with 0.5 mL of TES and the procedure was repeated once. Add 465 μL of Blocking buffer, 25 μL of BSA, 10 μL of Yeast t-RNA to the beads with TES removed, and rotate the beads for 2 h at 4 °C. Place the beads on a magnetic separation device to remove the supernatant, add 0.5 mL of lysis buffer, invert and mix, remove the supernatant, repeat the above steps once, and finally add 40 μL of lysis Lysis buffer to be used. Add 1.1 mL Lysis buffer, 11 μL protease inhibitor, 5.5 μL RNase Inhibitor, and 11 μL DTT to the cell lysis precipitate after mixing well in advance. The cells were placed on ice and lysed for 30 min, during which the cells were vortexed three times. 13000×g, centrifuged at 4 °C for 5 min, the supernatant was taken into a new EP tube, 1 mL of lysate was kept for subsequent experiments, and 100 μL was frozen as input at -80°C. Add the closed magnetic beads to the lysate and incubate for 4 h at 4 °C. After removing the supernatant, add the washing buffer and wash 5 times for 5 min each time. Elute RNA and precipitate RNA as in the experimental protocol of RAP, and follow up with reverse transcription and qPCR as described previously.

**Dual luciferase assay**

The pmiRGLO-circNP37 plasmid was constructed by GUANGZHOU IGE BIOTECHNOLOGY LTD. The interaction seed region of miR-361-5p/miR-345-5p with circNP37 was analyzed based on the predicted results of RNAhybird and miRanda. The pmiRGLO-circNP37 plasmid with seed region mutation was constructed as described in the Point mutation. 293T cells were cultured in 24-well plates the day before transfection experiments. Experimental treatment groups included: miR-361-5p+pmiRGLO-circNP37-WT, miR-361-5p+pmiRGLO-circNP37-Mut, NC-miRNA+pmiRGLO-circNP37-WT, NC-miRNA+ pmiRGLO-circNP37-Mut, three replicate were prepared for each treatment.

About 80% of the cells were confluent and prepared for plasmid transfection. Lipofectamine™ 3000 Reagent (Invitrogen™) was used in transfection experiment. Transfection complexes were prepared according to the instructions.

Samples were recovered 48 h post-transfection for fluorometric assay using Dual Luciferase Reporter Assay Kit (Vazyme). The cell supernatant in the culture dish was discarded, PBS was added and washed twice, and 100 μL of cell lysis solution was added to each well and lysed for 20-30 min. The cell lysates were centrifuged at 12,000 rpm for 2 min, 20 μL of supernatant was placed in a white opaque 96-well plate, 100 μL of Luciferase substrate balanced to room temperature was added to the cell lysates using a multichannel pipette, and the plate was tapped for immediate fluorescence measurement. Again using a multichannel pipette, add 100 μL Renilla substrate, tap the plate, and immediately perform the fluorescence measurement. For data calculation, the relative quantification method was chosen, and the fluorescence of each replicate well was used as one replicate, and the fluorescence intensity of Renilla Luciferase was used as the reference, and the NC-mimics treatment group was used as the experimental reference for normalization.

**Prediction of competitive endogenous RNA interactions**

Download all mature human miRNA sequences in 2022 from miRbase (https://www.mirbase.org/). Enter the circNP37 sequence and human mature miRNA sequences (both in Fasta format) in miRanda (http://www.bioinformatics.com.cn/local_miranda_miRNA_target_prediction_120) and the resulting predictions are saved in txt format. The experiments were performed using the default options. Enter the circNP37 sequence and the human mature miRNA sequence (both in Fasta format) in miRDB (https://mirdb.org/mirdb/index.html) and save the predicted results in txt format. The experiment is performed using the default options. Use Venn diagram (http://www.bioinformatics.com.cn/static/others/jvenn/example.html) to take the intersection of miRDB and miRanda prediction results. The binding strength of circNP37 to miRNA was calculated using RNAhybird (https://bibiserv.cebitec.uni-bielefeld.de/rnahybrid) with options energy Threshold = -10, approximate p-value = 3utr_human. The candidate miRNAs were also ranked from highest to lowest according to the absolute value of binding free energy of circRNA to miRNA and the number of binding sites. The candidate miRNAs were analyzed in miRmine (https://guanfiles.dcmb.med.umich.edu/mirmine/index.html) for whether they were expressed in lung and whether they could be expressed in A549/293T cells. The above experimental results were combined to summarize the candidate miRNAs of this experiment.

**Western Blot**

The cell supernatant was first removed from the six-well plate and the cells were gently washed twice with ice-cold PBS, and the experiment was performed on ice. Add 150 μL of RIPA lysate (Beyotime) containing 2% PMSF (Beyotime) directly to the well plate and place on ice for 3 min. Collect the cell lysate in an EP tube and place in a centrifuge at 4 °C for 5 min at 13,000×g. Add 40 μL of supernatant to 10 μL of 5 × Loading Buffer (Beyotime). After mixing, the samples were placed at 100°C for 5 min. PAGE Gel was prepared using the One-Step PAGE Gel Fast Preparation Kit (10%) (Vazyme). Insert the gel into the electrophoresis tank, carefully pull out the comb and add 20 μL of protein sample. 150 V electrophoresis for 60 min. The PVDF membranes were cut to 8.5 × 5.5 cm size and placed in methanol for 1-2 min of activation. The transfer using Rapid Transfer Solution (Beyotime) at 400 mA for 25 min. The PVDF membrane was gently washed once at the end of the transfer using TBST (prepared by diluting 10×TBS (Beyotime) to working concentration). The PVDF membranes were placed in 5% skim milk powder (Beijing Dingguo) TBST and in a horizontal shaker for 1 h at room temperature. After blocking, the PVDF membrane was gently washed once with TBST, and the membrane was cut according to the protein distribution using the Marker (Vazyme) as a reference. Place the membrane in an antibody incubation kit, add the working concentration of primary antibody and incubate at 4 °C for 16-19 h. After the primary antibody was recovered, TBST was added and the membrane was washed three times for 10 min each. Add the secondary antibody and incubate at room temperature with shaking for 1 h. After discarding the secondary antibody, add TBST and wash the membrane three times for 5 min each. Chemiluminescence Substrate (Millipore) was prepared, Chemiluminescence Substrate was added on the membrane, and the blot was exposed after 1 min of reaction, and the experimental results were counted for analysis. Grayscale analysis of protein bands was performed by ImageJ.

**In vivo experiments**

Five- to seven-week-old SPF BALB/C female mice were acquired and randomly divided into MOCK (n=7), WT-PR8-infected (n=10), and Mut-PR8-infected (n=11) groups. Mice were weighed and recorded prior to anesthetic injection. By intraperitoneal injection, each mouse was injected with 350 μL of Afodin (Afodin preparation: $ddH_2O$ was heated to 50°C, 0.25 g of tribromoethanol was first dissolved in 0.5 mL of 2-butanol, added to 20 mL of $ddH_2O$, filtered through a 0.22 μm filter and stored at 4 °C). After waiting for the mice to be deeply anesthetized, the virus diluted in PBS was infected by nasal drops. After infection, the behavior and weight change of the mice were recorded every 24 h. On the third day after infection, three to four mice from each group were randomly selected for dissection. Pathological sections were prepared from the left lung of the dissected mice, and the right lung was homogenized and used for the determination of murine lung viral load and viral titer. Mice lung pathology sections and H&E staining were completed by Servicebio. For, pathology sections, was scored by an analyst blinded to the experimental groups.

**Statistical analysis**

The two-sided Student's *t* test was used to compare the effects of two different treatments in parallel, after confirming that the data were normally distributed. $p < 0.05$ was considered statistically significant. $^*p < 0.05$, $^{**}p < 0.01$, $^{***}p < 0.001$. Statistical analysis was performed using SPSS. Histograms were plotted using GraphPad Prism 8 software. For in vivo experiments, changes in survival curves of mice were plotted using GraphPad Prism 8 software and analyzed for statistically significant differences.
